# Arbuscular mycorrhizal fungal families and exploration-based guilds exhibit distinct responses to N, P and K deficiencies and imbalances

**DOI:** 10.1101/2024.11.06.622229

**Authors:** Kian Jenab, Lauren Alteio, Ksenia Guseva, Stefan Gorka, Sean Darcy, Lucia Fuchslueger, Alberto Canarini, Victoria Martin, Julia Wiesenbauer, Felix Spiegel, Bruna Imai, Hannes Schmidt, Karin Hage-Ahmed, Erich M. Pötsch, Andreas Richter, Jan Jansa, Christina Kaiser

## Abstract

- Many agroecosystems face nitrogen (N), phosphorus (P) or potassium (K) deficiencies due to imbalanced or insufficient nutrient replenishment after plant biomass harvest. How this affects the symbiosis between plants and arbuscular mycorrhizal fungi (AMF), and the abundance of exploration-based AMF guilds (i.e., rhizophilic, edaphophilic, ancestral) remains largely unknown.
- We studied a 70-year nutrient-deficiency experiment in a managed grassland in central Austria, where aboveground biomass was harvested three times annually. N, P and K were fully, partially, or not replenished, causing long-term nutrient deficiencies and imbalances. We analysed AMF communities in soil and roots by DNA/RNA amplicon sequencing and fatty-acid biomarkers, alongside soil and plant community properties.
- Soil AMF communities were affected by N and P deficiencies, while root AMF communities were most susceptible to K deficiency, showing a 50% biomass reduction. We observed distinct guild- and family-specific responses: The edaphophilic guild declined with N deficiency, while the rhizophilic guild decreased with P and K deficiencies. Families within each guild, particularly in the ancestral guild, showed differential responses, indicating complementary nutrient specializations at the family level.
- Our findings underscore the previously unrecognized role of K deficiency in AMF symbiosis and suggest the existence of nutrient-related functional subgroups within exploration-based AMF guilds.

## Introduction

Arbuscular mycorrhizal fungi (AMF) are among the oldest and most important plant root symbionts (Sochorová *et al*., 2016; Thirkell *et al*., 2016; Brundrett & Tedersoo, 2018; Hawkins *et al*., 2023), forming associations with most plant species in croplands and managed grasslands. They provide essential soil nutrients like nitrogen (N), phosphorus (P), and potassium (K) to host plants (Smith & Smith, 2011; Berruti *et al*., 2016; Bravo *et al*., 2017; Igiehon & Babalola, 2017), along with other benefits such as pathogen protection (Borowicz, 2001), and soil structure improvement by aggregation (Rillig, 2004; Rillig & Mummey, 2006).

AMF are known to be sensitive to increasing soil nutrient availability from fertilization (Johnson *et al*., 2015). However, their responses to nutrient deficiencies and imbalances remain unclear. Although many agricultural soils are overfertilized in industrial countries, a substantial proportion of global agricultural soils suffer severe deficiencies of essential nutrients due to inadequate fertilization that fails to replace nutrients lost through crop harvest (Manning, 2012; Ludemann *et al*., 2024). Estimates indicate that, as of the year 2000, agricultural soils worldwide faced nutrient deficiencies driven by net losses of about 18.7 N, 5.1 P and 38.8 K (kg ha^-1^ yr^-1^) on 59%, 85% and 90% of harvested area, respectively (Tan *et al*., 2005). In particular, there is an increasing imbalance between K and other nutrients, driven by insufficient K replenishment compared to N and P in about 20% of global agroecosystems, particularly in Africa and China (Tan *et al*., 2012; Manning, 2012; Zörb *et al*., 2014). This problem has worsened with rising K fertilizer costs (Brownlie *et al*., 2024). Additionally, rising atmospheric N deposition due to anthropogenic activities increase N:P and N:K imbalances across all ecosystems, thereby fundamentally altering soil nutrient stoichiometry (Peñuelas *et al*., 2013; Lu & Tian, 2017).

Under low nutrient availabilities, AMF are essential partners for plants as their extraradical hyphal networks access inorganic nutrients (in particular P) more efficiently than plant roots (Smith & Smith, 2011). It is assumed that plants reduce support for AMF when nutrients are sufficiently available for efficient root uptake (Hoeksema *et al*., 2010). For example, high P fertilization commonly reduces AMF colonization in roots (Beauregard *et al*., 2010; Cavagnaro, 2015; Propster & Johnson, 2015; Liu *et al*., 2016). However, not only absolute quantities but also relative proportions (stoichiometry) of available nutrients are decisive (Johnson, 2010; Smith & Smith, 2011; Johnson *et al*., 2015): AMF biomass tend to respond positively to N fertilization under low P availability, but negatively when soil P availability exceeds a threshold (Johnson *et al*., 2015; Jiang *et al*., 2018; Han *et al*., 2020). In contrast to inorganic fertilizers, organic fertilizers often been enhance AMF growth, possibly due to slow nutrient release (Cavagnaro, 2015; Yang *et al*., 2018). Despite its importance, almost nothing is known about AMF responses to absolute or relative changes in K availability.

In addition to direct effects of nutrient deficiencies and imbalances on AMF, indirect effects via ecosystem feedback become important long term. For example, changes in vegetation or edaphic properties (e.g., pH, water or organic matter content (Jamiołkowska *et al*., 2018)) induced by nutrient deficiencies will additionally affect AMF. Particularly, changes in soil pH play an important role in AMF distribution and abundance (Clark, 1997; Van Aarle *et al*., 2002; Van Geel *et al*., 2018; Davison *et al*., 2021). At the same time, AMF communities influence ecosystem properties by shaping plant communities through changes in plant competitiveness, influencing soil physical structure, and building soil C stocks via their mycelia (Rillig & Mummey, 2006; Wang *et al*., 2016; Hawkins *et al*., 2023). The proportion and extent of AMF mycelia reaching into the soil - a trait of potential ecosystem relevance - is considered phylogenetically conserved at the family level (Hart & Reader, 2002; Powell *et al*., 2009). AMF families have thus been classified into ‘guilds’ based on exploration traits (Phillips *et al*., 2019; Weber *et al*., 2019). The ‘edaphophilic guild’ consists of families with extensive extraradical (i.e. outside plant roots in the soil) mycelia but limited intraradical (i.e. inside plant roots) mycelia, like Gigasporaceae. In contrast, the ‘rhizophilic guild’ includes families with large intraradical but sparse extraradical mycelium, such as Glomeraceae. The ‘ancestral guild’ comprises families with low biomass both intraradically and extraradically (Maherali & Klironomos, 2007; Weber *et al*., 2019). Each guild is associated with distinct functional traits, derived from mycelial structure and specific taxon- and family-level observations (Hart & Reader, 2002; Maherali & Klironomos, 2007; Treseder *et al*., 2018; Weber *et al*., 2019). Edaphophilic taxa are thought to be more efficient in supporting plant nutrient uptake and exhibit higher soil C sink strength due to their larger soil-based mycelium (Chagnon *et al*., 2013), while rhizophilic taxa are believed to be more effective in pathogen defense and conferring stress-tolerance (Chagnon *et al*., 2013; Weber *et al*., 2019), though some studies show members of Gigasporaceae may outperform Glomeraceae under biotic stress (Marro *et al*., 2022). Due to their properties, shifts in AMF guild composition under nutrient deficiencies and imbalances may significantly impact ecosystem functions, such as plant nutrient acquisition and soil C sequestration.

Although most plants in agroecosystems associate with AMF, it remains unknown how AMF communities will respond to increasing nutrient deficiencies and imbalances. Only few studies investigated long-term effects of nutrient deficiencies on AMF abundance and community composition in managed grasslands or croplands (Qin *et al*., 2015; Heyburn *et al*., 2017; Zheng *et al*., 2018). The responses of AMF exploration-based functional guilds to nutrient deficiency and imbalances remain largely unexplored. Moreover, while most studies focused on the effect of P or N, AMF responses to K deficiency and imbalances remain poorly understood.

In this study, we investigate the impact of over 70 years of N, P, and K deficiencies and imbalances on AMF communities in a managed grassland. We asked if and how long-term deficiencies and imbalances of N, P and K influence AMF composition and abundance in soil and roots, and whether this manifests in changes in exploration trait-based functional guilds. We tested whether responses of AMF communities were linked to responses of plant communities and soil properties.

We addressed these questions in a long-term nutrient deficiency experiment launched in 1946 in a permanent grassland site in Austria. We analyzed the effects of single and multiple N, P and K deficiencies on AMF abundance and community composition. For investigating the effect of pH, we included plots that were regularly limed. We investigated the intra- and extraradical AMF mycelium by analyzing soil and root samples, alongside assessing exploration-based AMF functional guilds through molecular profiling. We examined plant community composition and soil abiotic factors to identify potential links between these parameters and changes in AMF communities. We hypothesized that soil nutrient deficiencies would increase the relative share of edaphophilic taxa at the expense of rhizophilic and ancestral groups. We also hypothesized that P deficiency would have the strongest effect on AMF communities, and that deficiencies of single nutrients, as well as imbalances among N, P, and K would affect AMF community compositions. We further hypothesized that AMF responses would be linked to responses in soil properties and plant communities.

## Materials and Methods

### Study site

The study was conducted at a long-term nutrient deficiency experimental grassland site of the Agricultural Research and Education Centre Raumberg-Gumpenstein in Admont, Styria, Austria (47°34’58’’N, 14°27’02’’ E; 635 m a.s.l.). This grassland field experiment was established in 1946 on a hay meadow, with a three-cut per year management regime since start. According to the WRB soil classification system, the soil is a Gleyic Fluvic Dystric Cambisol with no supplementary qualifiers (FAO, 2015). The long-term mean annual precipitation is 1227 mm, and the mean annual temperature is 6.8 °C.

### Nutrient deficiency experiment

The experiment includes 24 nutrient-fertilization treatments (4 replicates each; 2.9 m × 7.1 m) in a randomized block design, of which 14 treatments were selected for our study (Fig. **S1**). The treatments comprise variations of inorganic (P, N, and K) or organic (solid manure and liquid slurry) fertilisation. Furthermore, some plots receive lime alongside inorganic fertilizers.

Nitrogen is applied as calcium ammonium nitrate (NH_4_NO_3_ + CaCO_3_), at 80 kg N ha^-1^year^-1^, split between spring and after the first cut. P was applied as Thomasphosphate ((CaO)_5_ P_2_O_5_ SiO_2_) from 1946 to 1997, and as hyperphosphate (Ca(H_2_PO_4_)_2_) thereafter at 35 kg P ha^-1^ year^-^ _1_, while K fertilization as KCl at 100 kg K ha^-1^ year^-1^, both in autumn. Some variants received additionally mixed lime (CaCO_3_ + MgCO_3_) at 928 kg Ca ha^-1^ every third year in autumn (referred to as ‘lime treatments’). Solid manure (37% N, 12% P, 51% K) and liquid slurry (35% N, 1% P, 64% K) are added at 15 t ha^-1^ (in autumn) and 40 t ha^-1^ (split between spring and post-cut), respectively. While all three nutrients are provided in the fully fertilized treatment, one or more nutrients are missing in the other treatments. Consequently, while aboveground plant biomass and nutrients have been removed consistently, only part of the nutrients have been replenished by fertilization. Additionally, the unfertilized control treatment has never received any fertilizer while being mowed three times a year since 1946. This practice has led to different types of long-term nutrient deficiencies in this grassland.

### Sample collection

In July 2019, soil samples were collected from 55 plots, comprising 14 treatments with 4 replicate plots each, except for the ’lime only’ treatment, which was sampled from 3 replicate plots. The samples were taken to 9.5 cm depth using 2 cm diameter corers. For each plot, 8 to 11 randomly distributed cores were combined into an approximately 300 g composite sample. Samples were sieved to 2 mm, and roots separated from soil. Fine roots were washed and cleaned. Subsamples for DNA and RNA extraction from root and soil samples were frozen in liquid N_2_ and stored at -80 °C. After lyophilization, root samples were ground in a cryomill with liquid N_2_ and both soil and root samples were stored at -20 °C for further analysis.

### Soil characterization

Soil pH (H_2_O) was measured in 1:5 (w:v) soil slurry. Soil water content was measured gravimetrically by weighing soil before and after drying at 105 °C for 24 h. For soil total C and N content, soil aliquots were dried at 60 °C for 72 h, ground and measured by EA-IRMS (EA 1110; CE Instruments, Milan, Italy), coupled to a Finnigan MAT Delta Plus IRMS (ThermoFisher Scientific, Waltham, MA, USA).

Plant available inorganic phosphate was measured photometrically by the phosphomolybdate blue reaction (Schinner *et al*., 1996), in 0.5M NaHCO_3_ extracts, with or without prior alkaline persulfate oxidation to estimate total dissolved P and dissolved inorganic P (Doyle *et al*., 2004). Ammonium was quantified in 1 M KCl soil extracts spectrophotometrically by Berthelot reaction (Tecan Infinite M200 Fluorimeter, Grödig, Austria) (Kandeler & Gerber, 1988; Hood-Nowotny *et al*., 2010). Nitrate was measured in 1 M KCl extracts by the VCl_3_ reduction and Griess reaction (Miranda *et al*., 2001; Hood-Nowotny *et al*., 2010). Soil total free amino acids were determined fluorimetrically based on Jones *et al*. (2002). Total dissolved N and dissolved organic C were measured in 1 M KCl soil extracts by TOC/TN-analyzer (TOC-V CPH/TMN-1, Shimadzu, Japan), and dissolved organic N was calculated by subtracting ammonium and nitrate from total dissolved N.

Microbial C, N and P were measured using chloroform fumigation-extraction (Vance *et al*., 1987), with modifications. Four g of soil from each plot replicate was weighed in two containers. To the first set of containers, 30 mL of 1 M KCl solution (for C and N) or 0.5 M NaHCO_3_ (for P) was added and shaken for 1 h before filtering through ash-free cellulose filters. The extracts were kept frozen at −20 °C for later analyses. The second set was fumigated with chloroform for 48 h in darkness, then extracted similarly. Dissolved organic C, total dissolved N and P were measured in the extracts as described above. Microbial biomass C was calculated by dividing fumigated and non-fumigated sample differences by 0.45, and microbial biomass N by 0.54; no factor was used for microbial biomass P (McDonald *et al*., 1998).

### Plant community composition

The most recent measurements of plant community composition (2015) and biomass (2018) were used, as they are the closest available measurements to our sampling date, ensuring the highest possible relevance to our study. Although we cannot rule out that plant community composition may have slightly shifted within a few years, we still think that this dataset holds valuable information as plant community composition tends to remain relatively stable over decades; for example, a Central European dry grassland showed no directional change in species composition over 90 years, despite interannual fluctuations (Fischer *et al*., 2020).

The plant community composition was determined in 2015 using the modified method of Braun-Blanquet (1951) described in Peratoner and Pötsch (2019). The plant species were identified taxonomically following Fischer *et al*. (2008). Plots were harvested in 2018 using a motorized bar mower and green mass yield was determined in the field. Representative samples were taken from each plot for dry matter content and forage quality analysis using standardized methods by VDLUFA (1976) and ALVA (1983).

### Neutral fatty acid biomarkers as proxy for AMF biomass

Neutral fatty acids (NLFAs) in soil and root samples were evaluated by the procedure described in Gorka *et al*. (2023). Total lipids were extracted with chloroform, methanol, and 0.15 M citrate buffer (pH = 4; v:v:v = 1:2:0.8) (Bligh & Dyer, 1959). Neutral lipids were isolated by elution from 96-well silica plates (Strata SI-1 Silica, 55 μm, 70 Å, 50 mg/well, Phenomenex, Canada) with chloroform containing 2% ethanol (Buyer & Sasser, 2012; Drijber & Jeske, 2019; Gorka *et al*., 2023). Subsequently, NLFAs were released from the lipids and transmethylated by alkaline methanolysis. Methyl-nonadecanoate (10 µg sample^-1^) was added as an internal standard for quantification of fatty acids by GC (7890B Agilent Technologies, USA) coupled to a TOF-MS (Pegasus BT, LECO, USA). NLFA 16:1ω5 was identified by comparison of chromatographic retention times to external fatty acid methyl ester standards (37 Component FAME mix, and BAME CP mix, Supelco, Austria), and by comparison to the NIST library of mass spectra. We used NLFA 16:1ω5 as a proxy for soil and root AMF biomass (Kaiser *et al*., 2015).

### AMF amplicon sequencing

AMF community composition was assessed by 18S rRNA gene amplicon sequencing using both RNA (cDNA) and DNA following Bukovská *et al*. (2021), using WANDA (CAGCCGCGGTAATTCCAGCT) and AML2 (GAACCCAAACACTTTGGTTTCC) primers. For RNA-based amplicon sequencing, total nucleic acids were treated with DNAse and reverse-transcribed to cDNA using SuperScript™ III (Invitrogen).

We conducted RNA and DNA amplicon sequencing in soil samples, while only DNA amplicon sequencing in root samples. In soil samples, total nucleic acids were extracted by phenol-chloroform procedure (Angel *et al*., 2012), whereas Qiagen DNeasy Plant Mini Kit was used to extract DNA from root samples. The amplicons were dually indexed with Nextera XT indexesand sequenced on Illumina MiSeq 2 × 300 platform (Nilsson *et al*., 2019; Bukovská *et al*., 2021). We used SEED software and external software packages to process sequencing results (Větrovský & Baldrian, 2013; Jansa *et al*., 2020). A minimum of 20 bp overlap was considered to pair raw sequence reads, and sequences with average quality scores below 30 were eliminated (Bukovská *et al*., 2021). Primers and chimers were removed from sequences, and reads were clustered at a 97% similarity threshold into operational taxonomic units (OTUs). These OTUs were identified using the SILVA database, retaining only Glomeromycota OTUs (Bukovská *et al*., 2021). Relative abundances of OTUs per sample were merged at genus level. AMF families were categorized based on their known growth traits (hyphae allocation in soil and roots) into three main guilds: rhizophilic, edaphophilic, and ancestral (Parniske, 2008; Powell *et al*., 2009; Weber *et al*., 2019).

DNA and RNA amplicon sequencing showed a comparable pattern of soil AMF communities (Fig. **S2**). However, DNA amplicon sequencing yielded insufficient sequencing depths for AMF communities in soil samples, as WANDA primer also amplified tardigrades, nematodes, and other invertebrates in soil. Due to higher sequencing depth yield, we used RNA amplicon sequencing of the 18S rRNA gene to assess soil AMF communities.

### Statistical analysis

Statistical analyses were conducted in R (v4.3.3), and plots generated using the package “ggplot2” (v3.4.4) (Hadley, 2016). We tested if N, P and K additions affected AMF and plant biomass as well as environmental variables in a full factorial way using a three-way analysis of variance (ANOVA) on the source data after checking normality and variance homogeneity. For non-normally distributed data, we applied transformations (square root, logarithmic, or exponential) to achieve normality. To assess lime application effects on AMF biomass, we performed a two-way ANOVA for lime-treated plots with lime application and inorganic nutrient fertilization as factors. For plots receiving organic fertilizers, we conducted a two-way ANOVA with solid manure and liquid slurry applications as factors.

Multivariate analyses used the “Vegan” package (v2.6-2) (Oksanen *et al*., 2007). We analyzed long-term nutrient deficiency influences on soil and root AMF and plant communities by correspondence analysis (CA), suitable for unimodal species distributions in ecological datasets (Ter Braak, 1985), and across treatments. CA1 scores represented AMF and plant community composition, as this axis explained most variation. We applied Permutational multivariate analysis of variance (PERMANOVA) to assess treatment effects on community compositions. We assessed soil and root AMF relationship with plant community compositions using the Mantel test, based on Bray–Curtis dissimilarity matrices and Pearson’s correlation coefficient. Associations between AMF abundance and community composition, and environmental variables were evaluated using Pearson’s or Spearman’s correlation for normally and non-normally distributed variables, respectively. Canonical correspondence analysis (CCA) was used to examine the influence of environmental factors on AMF and plant community compositions.

## Results

### Long-term fertilization regimes altered soil nutrient contents and pH

Over 70 years without replenishment of N, P or K removed by biomass harvest has significantly altered soil organic and inorganic pools of these elements. Compared to fully fertilized control plots, long-term absence of N fertilization significantly reduced total dissolved N, dissolved inorganic N, Ammonium and Nitrate (*P*<0.001; Fig. **S3d**; Table **S1**). Absence of N fertilization also reduced plant biomass and increased soil pH and inorganic P (Figs **1**, **S3a,f**; Table **S1**). Long-term absence of P inputs significantly reduced all measured P pools (dissolved organic P, inorganic P and microbial P) (Fig. **S3e,f**; Table **S1**). It also decreased plant biomass and soil pH, but increased dissolved organic C, soil C content, and some N pools (soil N content, total dissolved N, total free amino acids, and Ammonium) (Figs **1**, **S3a**; Table **S1**). Lack of potassium fertilization significantly increased soil pH and water content (Fig. **S3a,b**; Table **S1**). While we did not measure soil K, we refer to Pavlů *et al*. (2022), who reported plant-available K, P, Ca and Mg from the same long-term nutrient deficient experiment. Their findings show K fertilization substantially increased plant-available K in soil at this site with the highest mean value in the K-only treatment (238.8 mg kg-1), compared to about 98 mg kg^-1^ in the non-fertilized control. However, K fertilization only significantly enhanced soil K levels when N and P fertilization were missing, as higher plant biomass production in N and P fertilized plots likely offset K accumulation in soils (Pavlů *et al*., 2022). Soil pH was strongly affected by inorganic fertilizer regimes, with the lowest pH values (around 4.7) in plots that received N but no P, and the highest pH (up to 5.5) in those that received P, but no K (Fig. **S3a**; Table **S1**).

**Figure 1.**
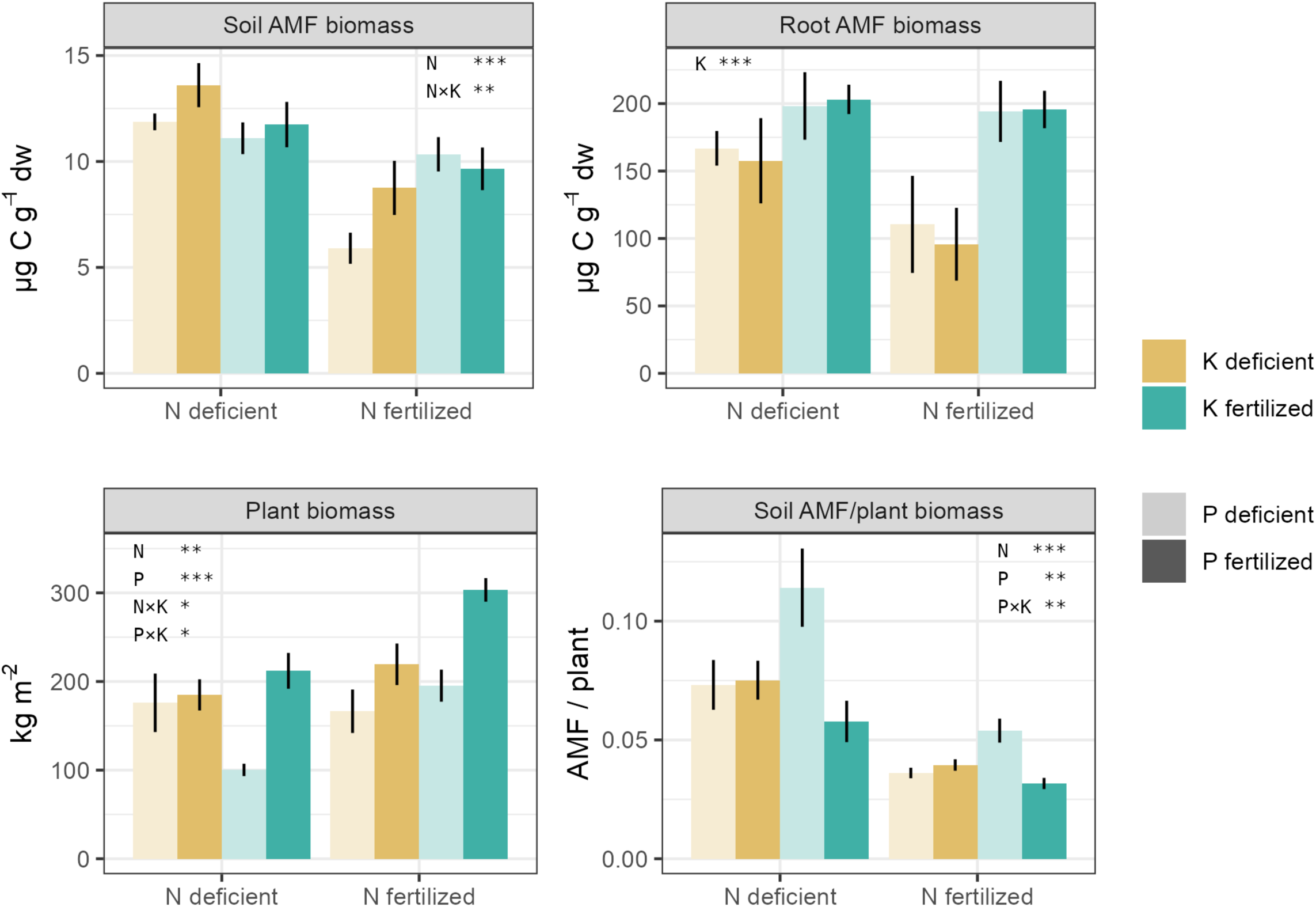
Long-term deficiencies of N, P and K affect AMF in soil and roots, and plant biomass. Soil AMF biomass increased with N deficiency, while root AMF biomass decreased with K deficiency (*P*<0.05, ANOVA, see Table 1). The panels show soil (i.e. extraradical) and root (i.e. intraradical) AMF biomass quantified using the neutral lipid fatty acid 16:1ω5 as a biomarker, as well as aboveground plant biomass and soil AMF biomass per plant biomass unit. X-axes depict N treatments (left bars: N-deficient, right bars: N-fertilized). Colours indicate K treatments (ochre: K-deficient, cyan: K-fertilized). Colour transparency indicates P treatments (light colors: P-deficient, dark colors: P-fertilized). Bars represent means ± standard error of the mean (n=4). Statistical significance from the three-way ANOVA is reported within each panel: *: *P* < 0.05, **: *P* < 0.01, ***: *P* < 0.001 (See also Table **1**)

**Table 1.**
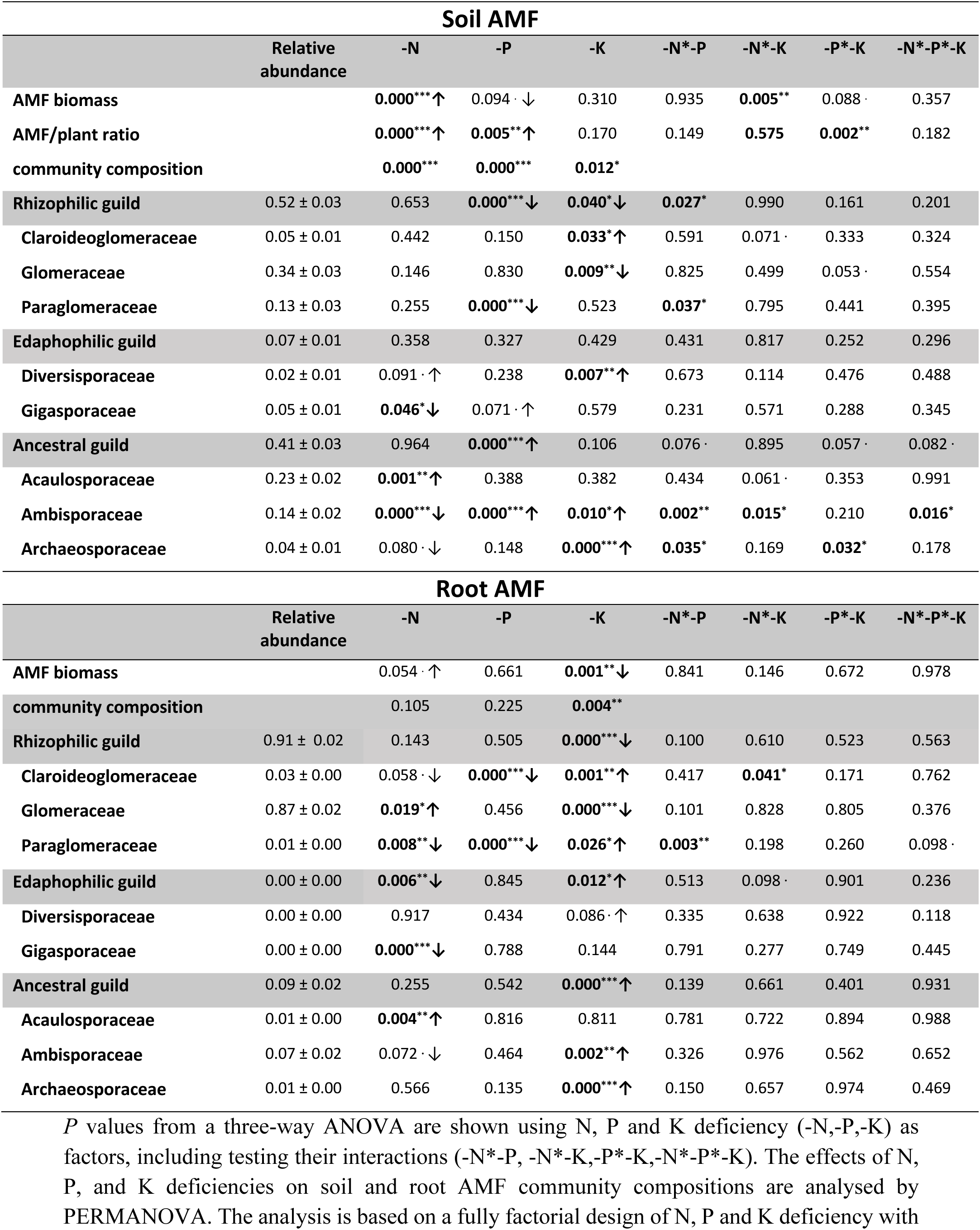
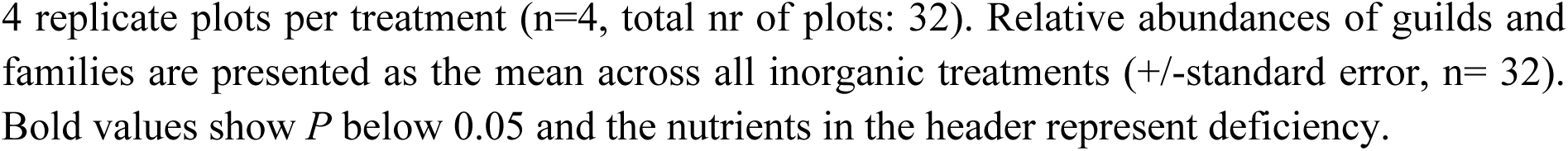
Effects of long-term N, P and K deficiencies on soil and root AMF biomass, families and guilds in inorganic treatments. *: *P* < 0.05, **: *P* < 0.01, ***: *P* < 0.001, . : *P* < 0.1, ↑: positive main effect, ↓: negative main effect *P* values from a three-way ANOVA are shown using N, P and K deficiency (-N,-P,-K) as factors, including testing their interactions (-N*-P, -N*-K,-P*-K,-N*-P*-K). The effects of N, P, and K deficiencies on soil and root AMF community compositions are analysed by PERMANOVA. The analysis is based on a fully factorial design of N, P and K deficiency with 4 replicate plots per treatment (n=4, total nr of plots: 32). Relative abundances of guilds and families are presented as the mean across all inorganic treatments (+/-standard error, n= 32). Bold values show *P* below 0.05 and the nutrients in the header represent deficiency.

### AMF biomass was linked to N and P in soils, but to K in roots

N, P and K deficiencies, caused by absence of respective fertilizer inputs, had strong and distinct effects on AMF biomass (using 16:1ω5 NLFA as a proxy) (Fig. **1**). The responses of AMF biomass in roots and soil (intra- and extraradical mycelia, respectively) were driven by different element deficiencies. While N deficiency significantly increased AMF biomass in soil (*F*=27.51, *P*<0.001), its effect on AMF biomass in roots was less pronounced (*F*=4.13, *P*=0.053; Fig. **1**; Table **1**). P deficiency had a weak negative influence on AMF biomass in soil (*F*=3.04, *P*=0.09), but no effect on AM fungi in roots (Fig. **1**; Table **1**). However, P deficiency significantly increased soil AMF biomass per unit of plant biomass, particularly under K fertilization (*F*=27.51, *P*=0.005; Fig. **1**; Table **1**). We also observed a significant interaction between N and K effects on soil AMF biomass (*F*=9.37, *P*<0.01; Table **1**), with lowest values in N-fertilized plots lacking both P and K (Fig. **1**). In contrast, K availability was the strongest predictor for root AMF biomass. K deficiency significantly diminished root AMF biomass (*F*=13.35, *P*<0.01; Fig. **1**; Table **1**).

Treatment effects on AMF biomass were also reflected in correlations of soil and root AMF biomass with soil N and P pools. Soil AMF biomass was negatively correlated with dissolved inorganic N pools (ammonium and nitrate), and the ratio of dissolved N to dissolved P, while root AMF biomass showed no correlation with soil N or P pools (but note that K pools were not measured in this study) (Fig. **2**).

**Figure 2.**
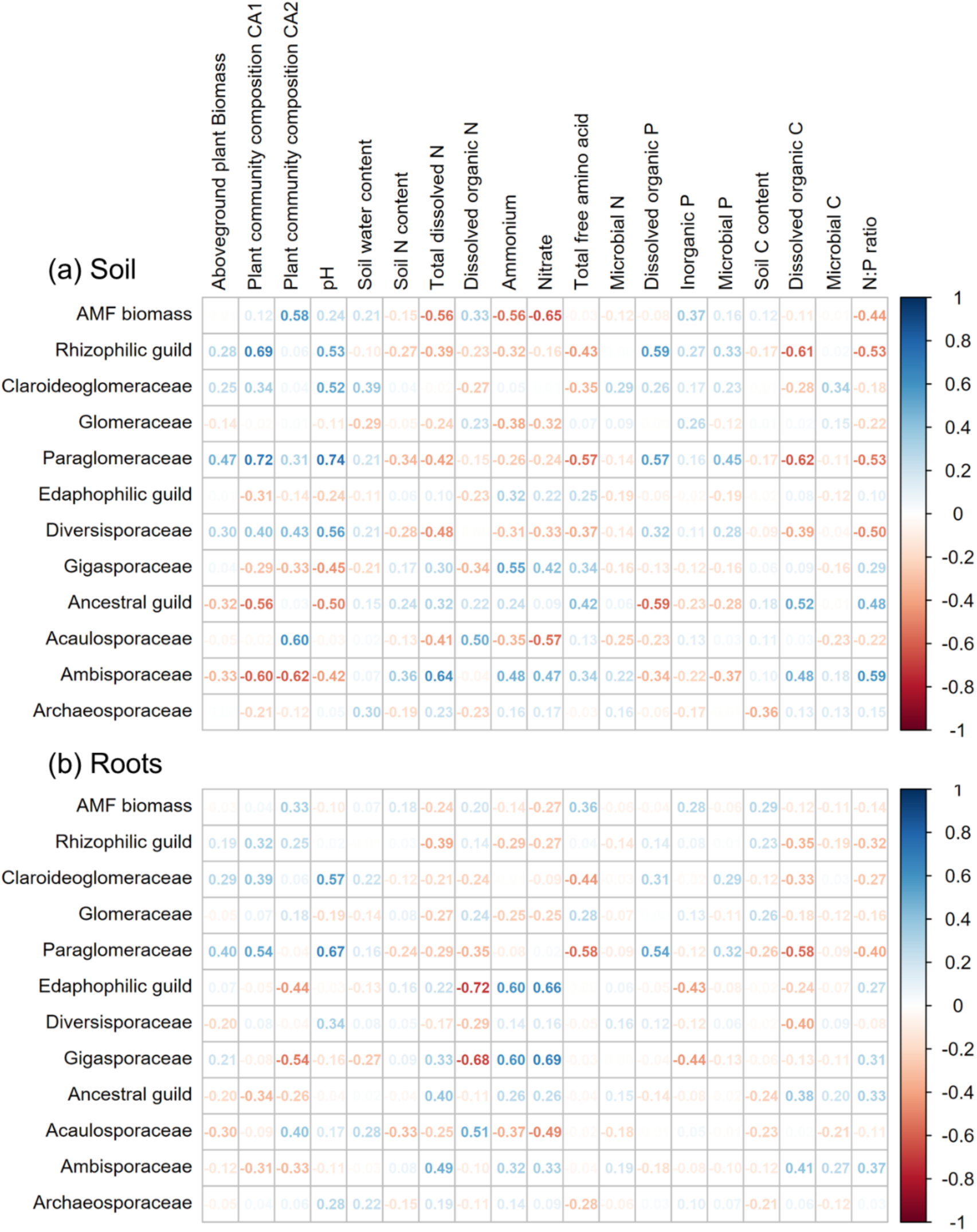
AMF biomass, guilds and families are correlated to plant and soil properties across N, P and K deficiency treatments. Numbers show pairwise Spearman’s correlation coefficients, with red and blue colors indicating negative and positive correlations, respectively. Correlation analysis was based on inorganically fertilized treatments only (including the unfertilized control). For correlation analysis among all treatments please see Fig. **S8**.

Organic fertilization increased AMF biomass in soils (but not in roots) compared to the fully fertilized (NPK) inorganic treatment (Fig. S**4**). Solid manure increased soil AMF biomass (*F*=6.11, *P*<0.05) but its combination with liquid slurry lessened this effect (*F*=8.50, *P*<0.05; Fig. **S4a**).

### AMF community composition differed systematically between soils and roots

In both soil and root samples, we found 11 AMF genera from eight families (Fig. **3**). Roots were dominated (approximately 75–99%) by AMF families and genera of the rhizophilic guild (mostly *Dominikia, Rhizophagus,* and *Glomus*; Fig. **3**). In contrast, soil samples contained substantial proportions (approximately 20–75%) of ancestral and edaphophilic taxa, particularly in inorganically fertilized plots (Fig. **3**).

**Figure 3.**
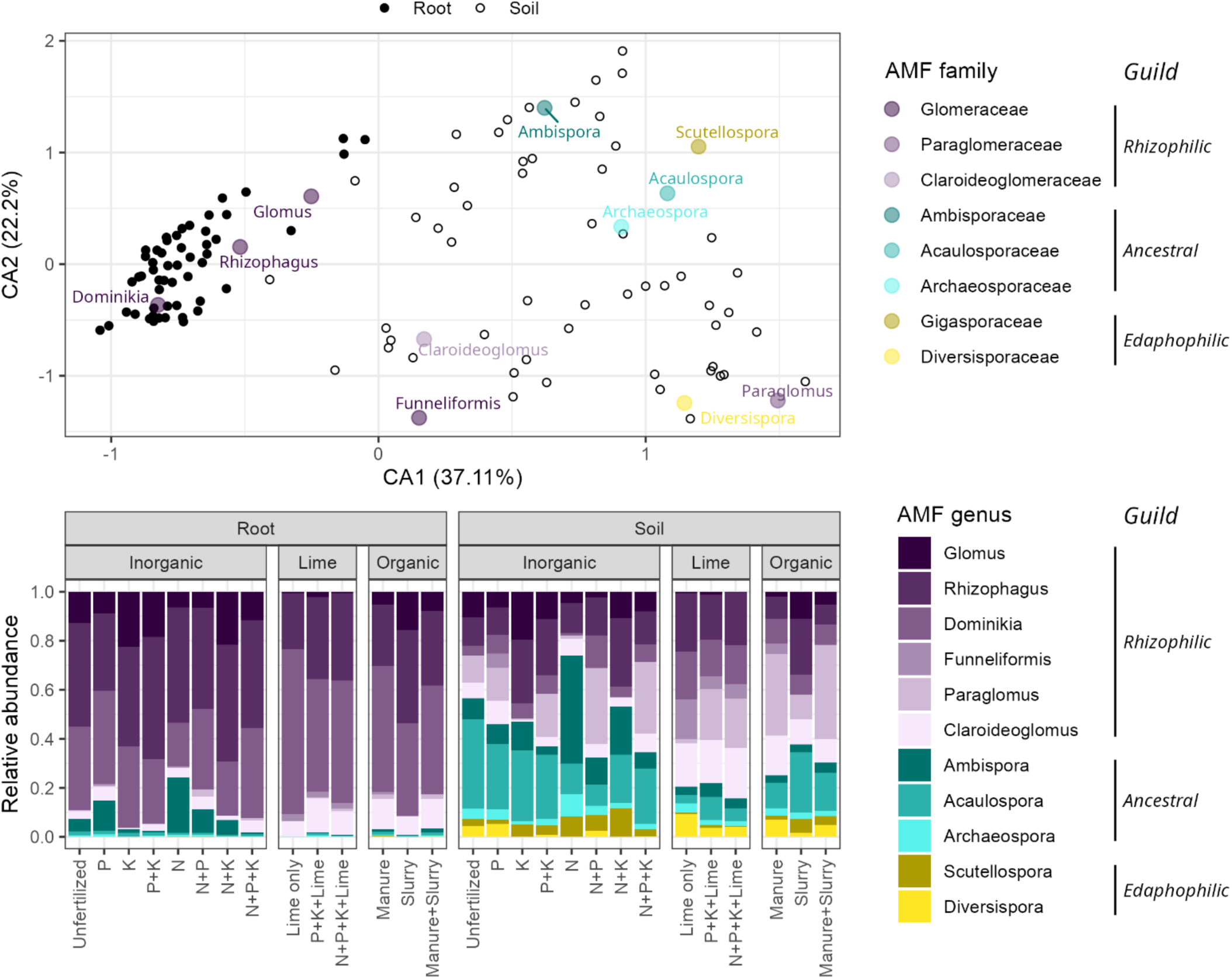
AMF community compositions differed systematically between soils and roots, and were shaped by long-term fertilization regimes. Root samples were significantly separated from soil samples in a multivariate correspondence analysis (CA) based on AMF genera (upper panel). Shown is a biplot of the first two CA axes, which together explain 59% of the total variability of the data. Small black-and-white circles represent soil and root samples (soil: open circles ○, root: closed circles ●). Larger colored circles indicate positions of AMF genera on the biplot, with colors indicating their families. Information on treatments associated with each data point in the CA biplot can be found in Fig. **S5**. The lower panel shows mean relative abundances of soil and root AMF genera for each long-term fertilization regime (n=4). The relative abundances of rhizophilic, ancestral, and edaphophilic guilds are depicted in purple, cyan, and yellow, respectively. Data shown in this figure is based on 18S rRNA gene amplicon sequencing using RNA (cDNA) for soil, and DNA for root samples (a comparison of soil DNA with root DNA data yielded similar results and can be found in Fig. **S2**).

Root samples were significantly separated from soil samples in a multivariate correspondence analysis (CA) over all treatments (PERMANOVA: *F*=81.35, *P*<0.001; Fig. **3**). *Rhizophagus*, *Dominikia*, and *Glomus* (all Glomeraceae) were more abundant in roots, while *Scutellospora* and *Acaulospora* (belonging to ancestral and edaphophilic guilds) were more abundant in soils. (Fig. **3**). This separation of root and soil samples was also visible when comparing soil and root DNA (Fig. **S2**), which rules out that it could have been caused by differences between DNA and RNA abundances in roots and soil, respectively.

### AMF community composition is associated with N and P in soils, but with K in roots

While the axis with the largest explanatory power separates root from soil samples (CA1, explaining 37.11% of the variability), the second axis (CA2, explaining 22.2% of the variability) is driven by the fertilization treatments (Figs **3**, **S5**). A clear separation exist among lime, organically fertilized, and the inorganically fertilized plots, which is consistent across root and soil AMF communities (PERMANOVA: *F*=8.90, *P*<0.001; Fig. **S5**). Additional multivariate analyses of inorganic treatments revealed different effects of nutrient deficiencies on AMF communities in roots and soils (Fig. **S5**). Soil AMF composition was mostly affected by N and P, and less by K deficiencies, while root AMF communities were significantly affected only by K deficiency (PERMANOVA analysis, Table **1**). This pattern, with N and P affecting soil AMF and K affecting root AMF communities, aligns with our AMF biomass findings (Fig. **1**; Table **1**).

### Distinct response of AMF guilds and families to inorganic nutrient deficiencies

We observed strong and clear responses of AMF families and guilds to nutrient deficiencies, which were mostly consistent across soil and roots. For example, P-deficient plots showed lower relative abundances of Paraglomeraceae (in roots and soil) and Claroideoglomeraceae (in roots), both belonging to the rhizophilic guild (*P*<0.001; Fig. **4**; Table **1**). At the guild level, the rhizophilic guild declined in P-deficient plots in soil, while the ancestral guild increased (Fig. **S7**), the latter mostly driven by an increase in Ambisporaceae (*P*<0.001; Fig. **4**; Tables **1**).

**Figure 4.**
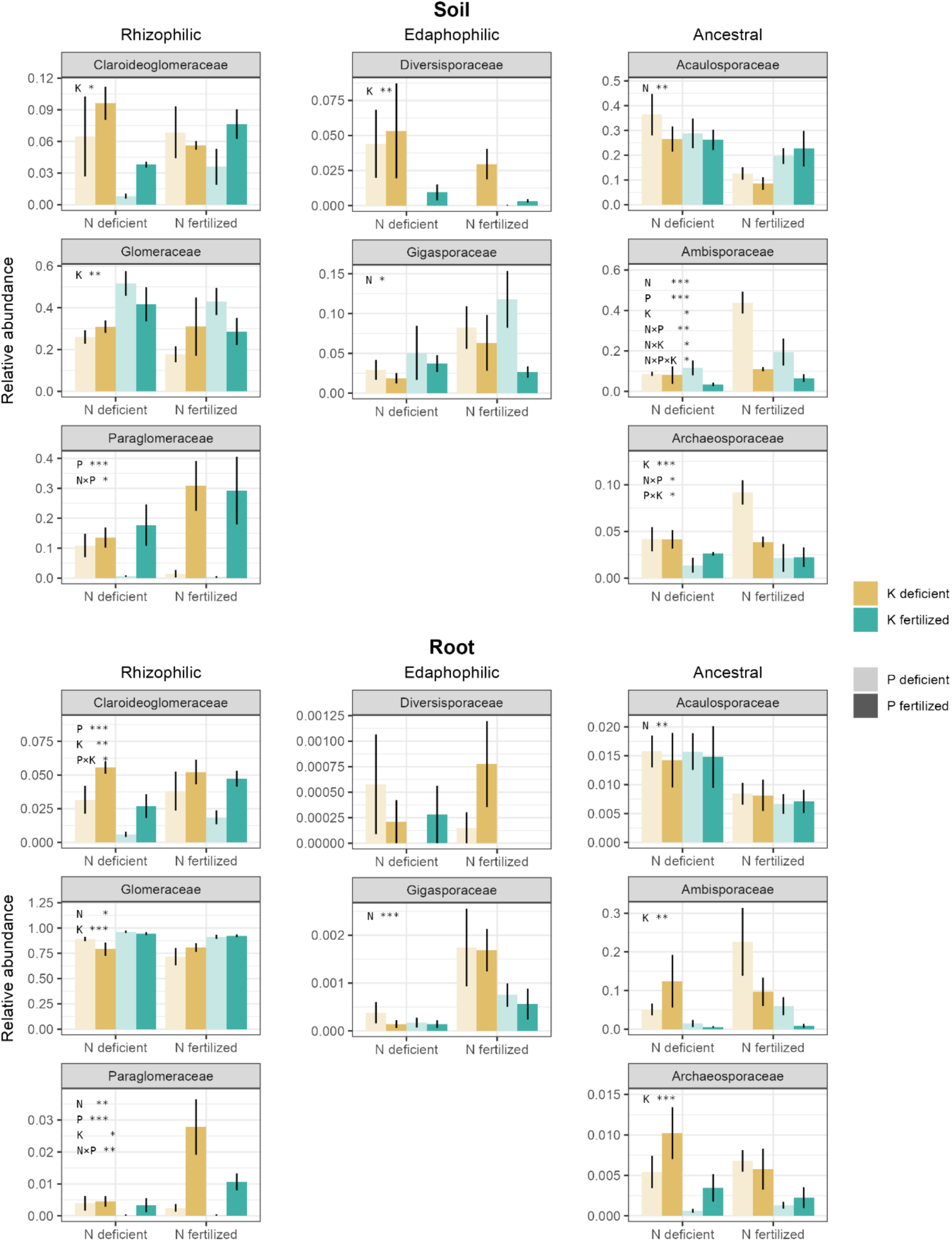
AMF families showed distinct responses to long-term deficiencies of N, P and K, which are partly consistent across soil (upper panel) and roots (lower panel). AMF families were grouped into exploration-based functional guilds (rhizophilic, edaphophilic and ancestral) according to Weber *et al*. (2019). Barplots show relative abundances of each AMF family on the total AMF. X-axes depict N treatments (left bars: N-deficient, right bars: N-fertilized). Colours indicate K treatments (ochre: K-deficient, cyan: K-fertilized). Colour transparency indicates P treatments (light colors: P-deficient, dark colors: P-fertilized). Bars represent means ± standard error of the mean (n=4). Statistical significance from three-way ANOVA is stated within each panel: *: *P* < 0.05, **: *P* < 0.01, ***: *P* < 0.001 (See Table **1**).

In contrast, N deficiency did not affect rhizophilic guild members in soil and roots, but decreased Gigasporaceae (edaphophilic guild) in both sample types, especially in roots (Figs **4**, **S7**; Table **1**). Moreover, N deficiency shifted families within the ancestral guild, increasing Acaulosporaceae and decreasing Ambisporaceae in soil, with similar trends in roots (Fig. **4**; Table **1**).

While P deficiency reduced less dominant rhizophilic families like Paraglomeraceae and Claroideoglomeraceae, K deficiency negatively affected the dominant rhizophilic family Glomeraceae in both roots and soils (Fig. **4**; Table **1**), shaping a significantly negative response of the rhizophilic guild to K deficiency (Fig. **S7;** Table **1**). Simultaneously, K deficiency increased ancestral families, mainly Archeasporaceae and Ambisporaceae in roots and soils, and edaphophilic guild in roots (Figs **4****, S7;** Table **1**).

The significant effect of long-term deficiencies and imbalances was also reflected in correlations between the relative abundance of families and soil N and P pools (Fig. **2**). For instance, Gigasporaceae positively correlated with ammonium and nitrate, while Acaulosporaceae showed negative and Ambisporaceae positive correlations with inorganic N. Paraglomeraceae correlated positively with organic P, while the ancestral guild was negatively associated with dissolved organic P in soil. Relationships between AMF families and soil N and P pools strenghtened when considering all treatments (Fig. **S8**).

### Changes in soil AMF communities were linked to changes in plant communities

Long-term nutrient deficiencies influenced plant communities. A correspondence analysis showed plant community composition was influenced by all three elemental deficiencies, with N and P deficiencies being more significant (PERMANOVA: *P*<0.001) than K deficiency (PERMANOVA: *P*<0.05, Fig. **S9**).

Plant community composition correlated significantly with soil AMF community composition, and less with root AMF community composition, as shown by a Mantel test (Figs **5**, **S10**). This was evident across all treatments (r=0.44, *P*<0.001 and r=0.31, *P*<0.001 for soil and root AMF, respectively; Fig. **S10**), but remained significant for soil AMF even in inorganic treatments (r=0.30, *P*<0.001 and r=0.08, *P*>0.05 for soil and root AMF, respectively; Fig. **5**)

**Figure 5.**
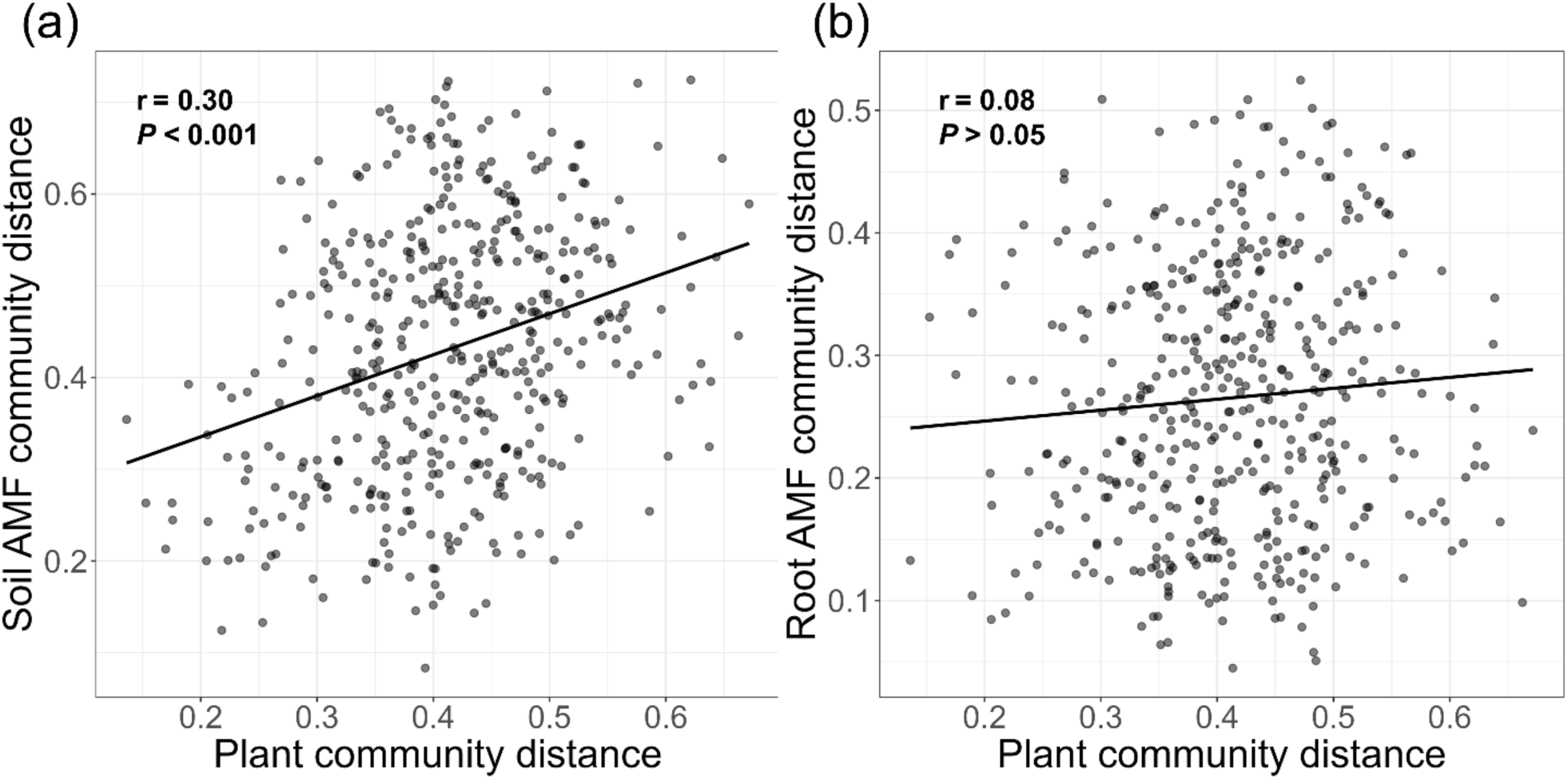
Stronger association of plant communities with soil than with root AMF communities. Mantel test results show correlations between a) plant and soil AMF communities, and b) plant and root-associated AMF community compositions across the inorganically fertilized treatments (including the unfertilized treatment). Each dot represents a pairwise comparison of community dissimilarities (Bray-Curtis distances) between plots. r indicates Pearson’s correlation coefficient between the distance matrices. For the association between soil and root AMF and plant communities by Mantel test among all treatments please see Fig. **S10**.

### Link between soil properties, and plant and AMF communities

We tested the link between soil properties and community composition of AMF and plants using a canonical correspondence analysis (Table **S2, S3**). pH and dissolved inorganic N correlated significantly with root and soil AMF communities in inorganic treatments (Table **S2**). Soil AMF communities were, as already indicated by the Mantel test, additionally linked to plant community composition (using the first correspondence axis of plant CA as a predictor; Table **S2**, Fig. **S9**). Like AMF communities, plant community composition was most linked to pH and dissolved inorganic N, but also to inorganic P. When all treatments were considered, these relationships remained, and additional predictors: dissolved organic N, P, and aboveground biomass for soil AMF; total free amino acids for root AMF; and soil total N for plants (Table **S3**) pH correlated strongly with the relative abundances of AMF guilds and families (Fig. **2**). It positively correlated with rhizophilic AMF, particularly Paraglomeraceae in soils and roots. In contrast, pH negatively correlated with the ancestal guild (particularly Ambisporaceae) and the family Gigasporaceae (Edaphophilic guild) in soil samples (Fig. **2**).

Notably, the strong correlation between pH and AMF community composition was also evident within inorganically fertilized treatments, excluding limed plots (Fig. **2**). Inorganic fertilizer regimes significantly affected pH, which ranged from 4.5 (plots that received N, but no P) to 5.5 (plots that received P but no K) (Fig. **S3a**; Table **S1**). Still, when considering all plots, we also observed a significant positive relationship between pH and AMF biomass (Fig. **S11b,d**). Liming substantially increased soil pH values (up to 6.5) and soil AMF abundance (*F*=86.33, *P*<0.001), especially in limed, but unfertilized plots where AMF biomass almost doubled (Fig. **S11a**). The positive effect of liming on soil AMF decreased when PK or NPK fertilizers were added (*P*<0.01 for both PK+Lime and NPK+Lime compared to PK and NPK treatments, respectively). The effect of liming on root AMF was smaller, but still significant (*F*=6.90, *P*<0.05; Fig. **S11c**).

## Discussion

Prolonged soil nutrient deficiencies disrupt the balance among plants, their AMF partners, and soil nutrient states. Predicting the outcomes of such disruptions is challenging due to the complexity of such interactions. Our findings show that 70 years of nutrient removal through biomass harvests significantly affected AMF communities in managed grasslands. Distinct fungal families responding differently to specific nutrient deficiencies, and these community shifts were linked to changes in plant composition and soil properties.

### Strong interplay between N, P and K determines soil AMF abundance and fungi-to-plant biomass ratios

AMF are considered crucial for P nutrition of grassland plants, while less is known on their role in N nutrition (Smith & Smith, 2011). We thus hypothesised that P availability has a greater impact on AMF communities than N. Contrary to this, N played the most important role for soil AMF biomass, with more AMF present in N-deficient compared to N-fertilized soils. P deficiency showed an opposing trend by decreasing soil AMF biomass (*P*=0.09), but only when K was also deficient. Both N and P deficiencies significantly increased the ratio of soil AMF- to-plant biomass, but only when K had been provided (Fig. **1**). This increase was driven by reduced plant biomass under these conditions (N or P deficiency with K fertilization), while AMF biomass remained stable (compare cyan bars in Fig. **1**). In contrast, in K-deficient plots, plant biomass remained relatively unaffected by additional P or N deficiency, while the negative effects of N fertilization and P deficiency on soil AMF biomass were enhanced. These results suggest that (i) plant P or N limitation in K-fertilized plots may have led to higher plant support of AMF per unit of (aboveground) plant biomass, as has been suggested for P limitation (Beauregard *et al*., 2010; Propster & Johnson, 2015; Hu *et al*., 2019) and (ii) this support may have been disrupted under plant K limitation, making it harder for AMF to cope with N and P limitation (see discussion on the effect of K limitation on plant physiology below). Moreover, K has been shown to enhance plant P uptake via both root and mycorrhizal pathways (Han *et al*., 2023) and a strong link between P and K has been demonstrated in mycorrhizas, suggesting a K and P interaction during mycelial transport (Garcia & Zimmermann, 2014). K limitation may thus constrain plant P uptake through the mycorrhizal symbiosis, explaining the lack of response of the soil AMF/plant biomass ratio to P deficiency in K-limited soils (Fig. **1**).

Stoichiometry-based concepts, like ‘trade balance’ and ‘functional equilibrium’ models suggest AMF symbiosis functionality depends on the N-to-P ratio (Johnson, 2010). If P is not added alongside N, increased N availability leads to aggravation of P deficiency in plants. Plants would therefore allocate more C to fungal partners to improve P uptake. Conversely, when P is abundant, plants reduce C allocation to AMF in favor of aboveground structures. Thus, it has been suggested that AMF biomass respond positively to N fertilization in soils when P is low, but negatively when P is abundant (Johnson, 2010; Wipf *et al*., 2019). However, our results contradict this pattern, showing no direct interaction between N and P in soil AMF biomass or AMF/plant biomass ratio, but strong interactions between either N or P with K (Fig. **1**; Table **1**). In our experiment, P fertilization included alkaline amendments, whose effect on soil pH may have altered our results compared to studies adding only P. Still, such strong interactions of P or N fertilization with K availability have not been shown before, possibly because most studies did not investigate K alongside N and P. Our results demonstrate that K plays an important and possibly so far-overlooked role in shaping AMF responses to varying nutrient availabilities.

### Lack of K fertilization strongly decreased root colonisation by AMF

Unlike soil AMF, which were predominantly influenced by N, root AMF were mainly affected by K, with a notable decrease in biomass under K deficiency (Fig. **1**; Table **1**). Previous studies, though limited, support a positive link between K and AMF root colonization. For instance, *Rhizophagus irregularis* colonization increased in Goji (*Lycium barbarum)* under high K levels (Han *et al*., 2023). Likewise, Liu *et al*. (2019) reported that K fertilization enhanced AMF colonization in mycorrhizal tomato plants.

Potassium is essential for higher plants, impacting carbon and water transport (by facilitating plant transpiration via stomata regulation), photosynthesis, nutrient uptake and stress response (Egilla *et al*., 2005; Cakmak, 2005; Wang & Wu, 2013; Sardans & Peñuelas, 2015). Particularly, K facilitates saccharide transfer from leaf parenchymal cells to the phloem (Deeken *et al*., 2002; Zörb *et al*., 2014; Sardans & Peñuelas, 2021) and is key for supplying sink organs with photosynthates (Zörb *et al*., 2014). Potassium deficiency reduces photosynthetic CO_2_ fixation and hinders partitioning and utilization of photosynthates (Cakmak, 2005). The reduced photosynthetic capacity and impaired transport of photoassimilates to below-ground plant organs and symbionts could explain the 50% reduction in AMF biomass in roots of K-deficient plots, and the lack of positive AMF response to P deficiency.

Impaired water transport in plants may also help to explain the significantly increased soil water content we observed in K-deficient plots (Fig. **S3**; Table **S1**). Although soil moisture could have been influenced by many different factors, we did not find any correlation except to the K treatments. As water content was however not related to AMF biomass in soils or roots (Fig. **2**), we assume that enhanced water contents did not directly affect AMF biomass.

Potassium deficiency increases NADPH oxidase activity, leading to reactive oxygen species (ROS) accumulation in roots (Cakmak, 2005; Garcia *et al*., 2017). Although AMF have been found to reduce ROS accumulation in plant roots during a 6-week K deficiency experiment with Medicago truncatula plants (Garcia et al, 2017), a prolonged K deficiency of 70 years could elevate ROS levels in plant roots to a degree that damages plant cell membranes and AMF in close proximity, creating unfavourable conditions for colonisation.

Notably, the impact of K deficiency was strongest on Glomeraceae AMF, which dominated in roots (87% of root AMF) but were less abundant in soils (Fig. **4**; Table **1**). This might explain the more significant effect of K deficiency on root AMF versus soil AMF. While we cannot, with existing knowledge, explain why Glomeraceae would be more susceptible to K deficiency, we can speculate that soil-based AMF families may have benefited more from altered soil properties like increased soil pH or water content in K-deficient plots, while root-centered Glomeraceae, were more affected by disturbed plant physiological processes due to K deprivation.

### Proportions of rhizophilic, edaphophilic and ancestral guilds differed between root and soil samples

We found that the rhizophilic guild (particularly Glomeraceae) were relatively more abundant in roots, while ancestral and edaphophilic guilds predominated in soil (Fig. **3**). For edapho- and rhizophilic AMF, this reflects their exploration traits: rhizophilic AMF produce a relatively higher proportion of hyphae inside roots, while edaphophilic AMF produce relatively more extraradical hyphae (Powell *et al*., 2009; Weber *et al*., 2019). Although ancestral AMF are thought to produce low amounts of both extraradical and intraradical hyphae (Powell *et al*., 2009; Weber *et al*., 2019), we found them in surprisingly high proportions in both soil and roots, exceeding even edaphophilic AMF in soils (Fig. **3**). This could indicate that ancestral AMF produce more hyphae than previously thought, or suggest a higher number of ancestral than edaphophilic individuals. Interestingly, some ancestral families (Acaulosporaceae and Archaeosporaceae) and even one rhizophilic family (Paraglomeraceae) were relatively more abundant in soils than roots (Fig. **3**), suggesting they may exhibit more ‘edaphophilic’ exploration traits than commonly assumed. This, together with our findings about their specific response to nutrient limitations, indicates that ancestral AMF may possess unique functional traits and may play more important ecosystem roles than previously thought (Maherali & Klironomos, 2007).

### AMF functional groups and families show highly specific responses to nutrient deficiencies

Although root and soil AMF biomass responded differently to nutrient deficiencies, responses of individual AMF families and guilds remained consistent across both sample types (Table **1**). The specific and significant responses of individual AMF families to nutrient deficiencies suggest that soil nutrient conditions exerted strong selection pressure at the family level. Although we grouped AMF families into functional guilds based on exploration traits (Weber *et al*., 2019), contrasting responses within guilds suggest that additional functional traits at the family level shape responses to nutrient limitations.

While little is known about other AMF functional traits at family level, exploration traits have been used to explain family- and guild-level responses to nutrient availability (Powell *et al*., 2009; Treseder *et al*., 2018; Weber *et al*., 2019; Han *et al*., 2020). Some reviews proposed that fungi with high soil exploration capabilities, like Gigasporaceae, decline in N-enriched soils due to reduced plant C allocation (Treseder *et al*., 2018; Cotton, 2018; Lilleskov *et al*., 2019), while fungi with limited soil exploration capabilities, such as Glomeraceae, might be favored (Treseder *et al*., 2018), possibly due to increased plant pathogen pressure at high N availability (Sun *et al*., 2020). This contrasts with the trade-balance model, which predicts plant preference for high-exploration AMF under N enrichment to offset emerging relative P limitation (Johnson, 2010).

Our results oppose the concept that N fertilization reduces the fraction of edaphophilic taxa. Although fewer AMF could thrive in N-fertilized soil, the relative abundance of the edaphophilic guild (mostly Gigasporaceae) increased in soil and roots (Fig. **S7**, Table **1**). In line with a recent global-scale meta-analysis showing that N addition decreased Glomeraceae abundance while minimally affecting Gigasporaceae (Han *et al*., 2020), we observed a negative effect of N addition on Glomeraceae, although this was significant only in roots (Fig. **4**; Table **1**). Interestingly, the ancestral family Acaulosporaceae, which showed an ‘edaphophilic’-like hyphal distribution, responded positively to N deficiency, contrasting with Gigasporaceae. We have to note, however, that our amplicon sequencing-based data only shows relative shifts among AMF families. As N-deficiency increased total AMF biomass (reflected by NLFA 16:1ω5), absolute numbers of Gigasporaceae may have remained stable, while Acaulosporaceae increased both relatively and absolutely. The contrasting behaviour of Gigasporaceae and Acaulosporaceae is also demonstrated by Gigasporaceae correlating positively to inorganic N (ammonium and nitrate) and negatively to organic N, while Acaulosporaceae showed the opposite pattern (Fig. **2**). Why N deficiency favours Acaulosporaceae over Gigasporaceae remains unclear. We can only speculate that Acaulosporaceae are more effective at N aquisition when mineral N is limiting, while Gigasporaceae more effectively provide other nutrients, such as P when they become stoichiometrically limiting, as proposed by the trade-balance model (Johnson, 2010).

Contrary to N, which mostly affected edaphophilic and ancestral guilds, P and K primarily affected rhizophilic families (Fig. **4**; Table **1**). While K deficiency strongly decreased Glomeraceae, P deficiency significantly decreased Paraglomeraceae in both soil and roots, and Claroideoglomeraceae in roots. Correspondingly, Paraglomeraceae showed the strongest positive correlation with dissolved organic P compared to other families, followed by Claroideoglomeraceae (Fig. **2**). While previous studies revealed a positive link between P and Paraglomeraceae (Kurle & Pfleger, 1996; Hijri *et al*., 2006), others found no such association (Gosling *et al*., 2014). Both families (particularly Paraglomeraceae), however, showed the strongest correlation among all families with soil pH in soil and roots. As the P fertilizer contained alkaline additions, which increased soil pH, the increase in Paraglomeraceae and Claroideoglomeraceae could thus also reflect a preference for higher pH, as has often been found for AMF in general (Helgason & Fitter, 2009; Han *et al*., 2020).

The negative impact of P deficiency on Paraglomeraceae contrasted with its positive effect on Ambisporaceae (ancestral guild), which increased in P-deficient soils (Fig. **4**; Table **1**). Ambisporaceae were positively correlated with ammonium, nitrate and N:P ratio, negatively with dissolved organic P, and with pH (in soil only) (Fig. **2**). Together, our results suggest contrasting functional roles between certain rhizophilic (especially Paraglomeraceae) and ancestral (particularly Ambisporaceae) families regarding P availability. Notably, Paraglomeraceae and Ambisporaceae were strongly associated with plant community composition in opposite directions (Fig. **2**). This indicates that their switch with P deficiency was linked to plant community changes, suggesting a certain degree of host-preference. Interestingly, Ambisporaceae showed the strongest positive response to P deficiency, surpassing Gigasporaceae (Table **1**), a commonly studied family under nutrient limitation (Powell *et al*., 2009; Treseder *et al*., 2018). We hypothesize that Ambisporaceae might play a critical yet underappreciated role in P uptake from P-limited soils.

Similar to P deficiency, K deficiency shifts the community from rhizophilic families to ancestral and, to some extent, edaphophilic families. K deficiency decreased the dominant rhizophilic family, Glomeraceae, while increasing Archaeosporaceae and Ambisporaceae (ancestral guilds) and Diversisporaceae (edaphophilic guild) (Fig. **4**; Tables **1**). This shift likely reflects a plant strategy to rely on AMF with more extensive extraradical mycelia for P and K uptake under limiting conditions provided by ancestral and edaphophilic taxa. Like P, K is relatively immobile in soil due to strong adsorption to particles, leading to root depletion zones. AMF are thus essential for plant K uptake (Garcia & Zimmermann, 2014). Our results indicate that Archaeosporaceae, Ambisporaceae and Diversisporaceae might possess traits supporting plant K uptake under K-limiting conditions, or provide adaptation mechanisms to host plants under nutrient stress (Garcia *et al*., 2017).

Noteworthy, we observed contrasting family-level responses to specific nutrient deficiencies within each exploration-trait-based guild. While Ambisporaceae and, to a lesser extent, Gigasporaceae may support plants under P deficiency, Archaeasporaceae, Ambisporaceae and the Diversisporaceae seem to play a critical role under K deficiency. Notably, N deficiency uniquely increased Acaulosporaceae (ancestral guild). Our analysis indicates that families with specific and complementary nutrient specializations exist within each guild, particularly high nutrient deficiency competence within the ancestral guild, whose families often behaved antagonistically to families of other guilds. Investigating functional properties of understudied AMF families will help expand the concept of AMF functional groups and enable targeted exploration of plants with specific AMF to reduce fertilizer use.

### AMF were closely linked to plant communities and soil pH

AMF are not host-specific (Klironomos, 2000), but exhibit host preference (Vandenkoornhuyse *et al*., 2002; Martínez-García *et al*., 2015). Different plant functional groups select distinct AMF communities potentially based on fungal life-history traits (Chagnon *et al*., 2013; Davison *et al*., 2020; Blažková *et al*., 2021). While these factors may explain the link between plant and soil AMF communities, both could also have responded similarly to changing soil parameters. Research showed soil conditions were more influential in shaping AMF communities in European grasslands than host plant specificity (Van Geel *et al*., 2018). Edaphic factors may exert stronger co-structuring effects on the soil-dominant AMF proportion, as extraradical hyphae are more exposed to edaphic parameters than intraradical hyphae. This could explain the stronger correlations between plant and soil AMF communities than between plant and root AMF communities (Fig. **5**).

Among all edaphic factors, soil pH showed the strongest link to both plant and AMF communities (Tables **S2, S3**). AMF prefer neutral or alkaline soil pH (Helgason & Fitter, 2009; Han *et al*., 2020), explaining higher AMF biomass in limed plots (Fig. **S11**). The absence of liming kept pH within a moderately acidic range, possibly accounting for the lack of correlation between AMF biomass and pH in inorganic treatments (Fig. **2**). However, pH remained the best explanator for soil AMF community composition even across inorganic treatments (Table **S2),** confirming previous findings highlighting narrow ecological niches for AMF, particularly for pH (Kawahara *et al*., 2016; Davison *et al*., 2021). The strong correlations between soil pH and both AMF and plant community compositions suggest strong interdependencies.

Interestingly, while topsoil C content significantly varied with fertilization regimes (Table **S1**), it did not correlate with AMF families, guilds or biomass (Figs **2****, S8**), despite significant shifts across exploration-based guilds (f.e. reduction of rhizophilic guild from 70% to 24% in P- and K-codeficient plots, Fig **S7**). Although AMF guilds are thought to affect soil C stocks through different mycelial structure and biomass (Horsch *et al*., 2023), we conclude that this may not be pronounced in agricultural settings, where fertilization-driven changes in decomposition and fine root biomass likely overrule the effect of AMF functional group distribution.

## Conclusion

Our research shows that N, P, and K deficiencies caused by biomass harvest without adequate element replacement significantly alter AMF biomass and communities. Prolonged K deficiency, a relatively understudied factor, greatly affects AMF and modulates the N and P effects. Notably, the combination of N fertilisation with inadequate K-fertilisation, widespread in global agriculture (Tan *et al*., 2012; Manning, 2012; Zörb *et al*., 2014), may be most detrimental for AMF biomass and root colonisation, and cause strong shifts in AMF families and guilds, with yet unknown effects on ecosystem functioning.

We reveal specific and complementary nutrient specializations at the AMF family level within exploration-based guilds. Understanding these responses is key to improving agricultural AMF inoculation strategies, which focus primarily on Glomeraceae taxa (Zhang *et al*., 2019; Lutz *et al*., 2023). Our study highlights the need to better understand the diverse spectrum of AMF families, especially ancestral ones, which is only possible through molecular approaches that overcome traditional biases towards Glomeraceae.

These findings highlight the intricate role of K alongside N and P in shaping AMF community structure and function. Moving beyond model taxa, our results advocate for incorporating diverse AMF families with complementary nutrient specializations into agricultural management, fostering resilient and efficient symbioses that are able to address widespread nutrient imbalances.

## Acknowledgement

This work has been supported by the European Research Council under the European Union’s Horizon 2020 research and innovation programme (grant agreement No 819446 to CK) and by the Austrian Science Fund, Cluster of Excellence COE7. The contribution of JJ was supported by the project No CZ.02.01.01/00/22_008/0004597, funded by the Czech Ministry of Education, Youth and Sports.

## Author contributions

**KJ:** Conceptualization, Formal Analysis, Investigation, Visualization; **LA:** Formal Analysis; **KG:** Formal Analysis, Visualization, Methodology; **SG:** Formal Analysis, Investigation, Visualization; **SD:** Investigation; **LF:** Formal Analysis, Investigation; **AC:** Formal Analysis, Investigation; **VM:** Investigation; **JW:** Investigation; **FS:** Investigation; **BI:** Investigation; **HS:** Investigation; **KH:** Conceptualization; **EMP:** Funding Acquisition, Project Administration, Resources; **AR:** Conceptualization, Funding Acquisition, Project Administration, Supervision, Resources; **JJ:** Conceptualization, Formal Analysis, Investigation, Methodology, Supervision; **CK:** Conceptualization, Funding Acquisition, Project Administration, Supervision, Validation, **KJ** and **CK** wrote the original draft and all other co-authors contributed to review and editing.

## Data availability

Sequencing data are available through NCBI - GenBank under accession number PRJNA1140020. All other data can be found in a public repository: https://doi.org/10.5281/zenodo.13760726

## Supporting Information

**Fig. S1** Experimental plot setup.

**Fig. S2** The effects of long-term nutrient deficiencies on soil (DNA-based) and root (DNA-based) AMF communities.

**Fig. S3** The impact of long-term nutrient deficiencies on soil properties and nutrient content

**Fig. S4** Effects of long-term organic fertilization (Manure and slurry) on soil and root AMF biomass..

**Fig. S5** Compositions of AMF communities in combined soil and root samples exposed to different long-term nutrient deficiencies evaluated by correspondence analysis (CA).

**Fig. S6** The effects of long-term nutrient deficiencies on soil and root AMF community composition based on correspondence analysis (CA).

**Fig. S7** The effects of long-term deficiencies of N, P and K on soil and root AMF functional guilds.

**Fig. S8** Relationship between AMF and environmental parameters across all treatments.

**Fig. S9** The impacts of long-term nutrient deficiencies on plant community composition.

**Fig. S10** Relationship of plant communities with soil and root AMF communities across all treatments.

**Fig. S11** The impact of long-term lime application and fertilization regime on soil and root AMF biomass as well as the relationship of soil and root AMF biomass with pH.

**Fig. S12** Correlations between AMF genera and environmental factors.

**Table S1** Three-way ANOVA results regarding the effect of long-term deficiencies of N, P and K on edaphic factors in inorganic treatments.

**Table S2** Results of canonical correspondence analysis (CCA) testing the effect of environmental variables on soil and root AMF, and plant community compositions across the inorganic treatments.

**Table S3** Results of canonical correspondence analysis (CCA) testing the effect of environmental variables on soil and root AMF, and plant community compositions across all treatments.

