## Supplementary Material for "Arbuscular mycorrhizal fungal families and exploration-based guilds exhibit distinct responses to N, P and K deficiencies and imbalances"

Article title: Strong family- and guild-specific responses of arbuscular mycorrhizal fungi to long-term nutrient deficiencies and imbalances

Authors: Kian Jenab, Lauren Alteio, Ksenia Guseva, Stefan Gorka, Sean Darcy, Lucia Fuchslueger, Alberto Canarini, Victoria Martin, Julia Wiesenbauer, Felix Spiegel, Bruna Imai, Hannes Schmidt, Karin Hage-Ahmed, Erich M. Pötsch, Andreas Richter, Jan Jansa, and Christina Kaiser

The following Supporting Information is available for this article:

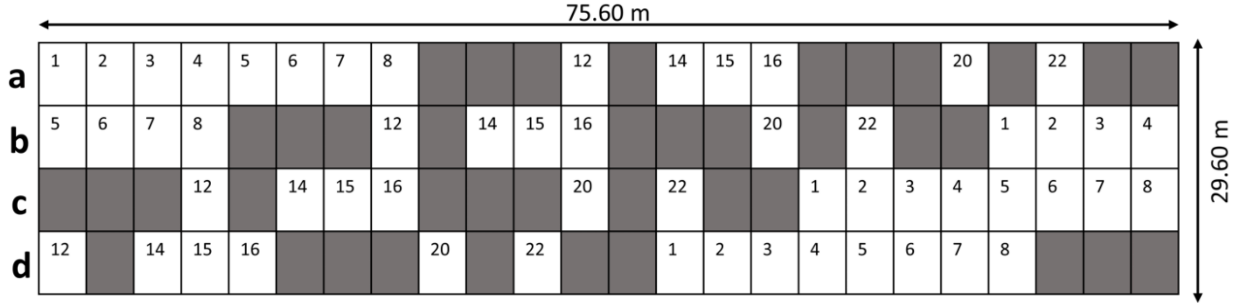


| Inorganic Treatment | Lime Treatment | Organic Treatment |
|---|---|---|
| Control (NPK fertilized) (12) | Control (NPK + Lime fertilized) (15) | Solid manure and liquid slurry (full fertilization) (22) |
| Full Deficiency (no fertilizer) (1) | Deficient (only lime fertilized) (2) | Solid manure (NP fertilized but with less N) (16) |
| P deficiency (NK fertilized) (6) | N deficient (PK + lime fertilized) (14) | Liquid slurry (mainly N) (20) |
| K deficiency (NP fertilized) (7) | | |
| N deficiency (PK fertilized) (8) | | |
| NK deficiency (P fertilized) (3) | | |
| NP deficiency (K fertilized) (4) | | |
| KP deficiency (N fertilized) (5) | | |

**Fig. S1: Experimental plot setup.** The figure indicates the spatial distribution of selected nutrient deficiency treatments for sampling at the Admont grassland experimental site. The numbers in the figure show different treatments and the detail about the treatments is explained in the table below the figure. Furthermore, the letters in the figure (a, b, c and d) show different replicate plots.

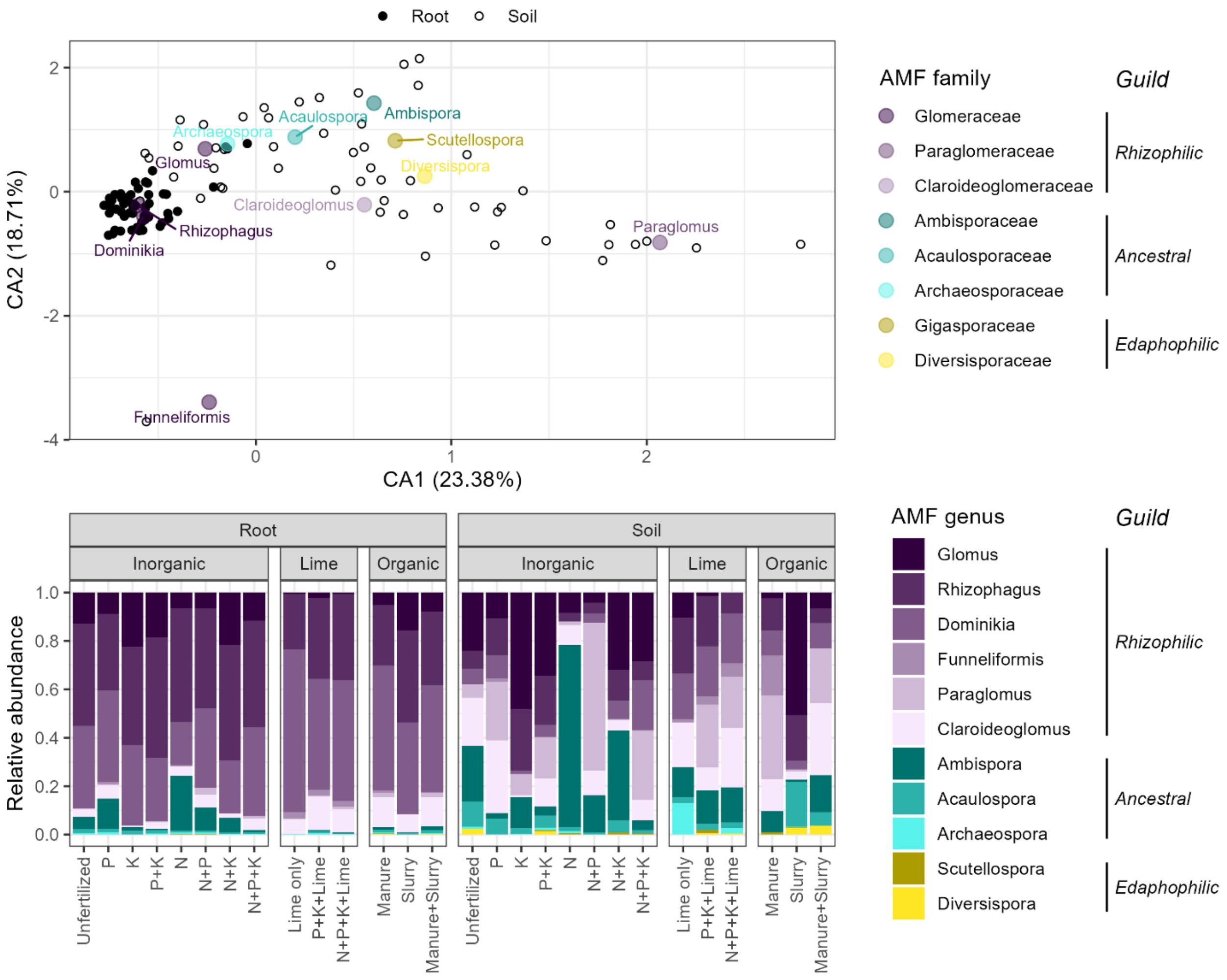


**Fig. S2:** AMF community compositions differed systematically between soils and roots, and were shaped by long-term fertilization regimes. Root samples were significantly separated from soil samples in a multivariate correspondence analysis (CA) based on AMF genera (upper panel). Shown is a biplot of the first two CA axes, which together explain 59% of the total variability of the data. Small black-and-white circles represent soil and root samples (soil: open circles ○, root: closed circles ●). Larger colored circles indicate positions of AMF genera on the biplot, with colors indicating their families. The lower panel shows mean relative abundances of soil and root AMF genera for each long-term fertilization regime (n=4). The relative abundances of rhizophilic, ancestral, and edaphophilic guilds are depicted in purple, cyan, and yellow, respectively. Data shown in this figure is based on 18S rRNA gene amplicon sequencing (DNA) for root and soil samples (a comparison of soil RNA with root DNA data yielded similar results and can be found in Fig. **3**).

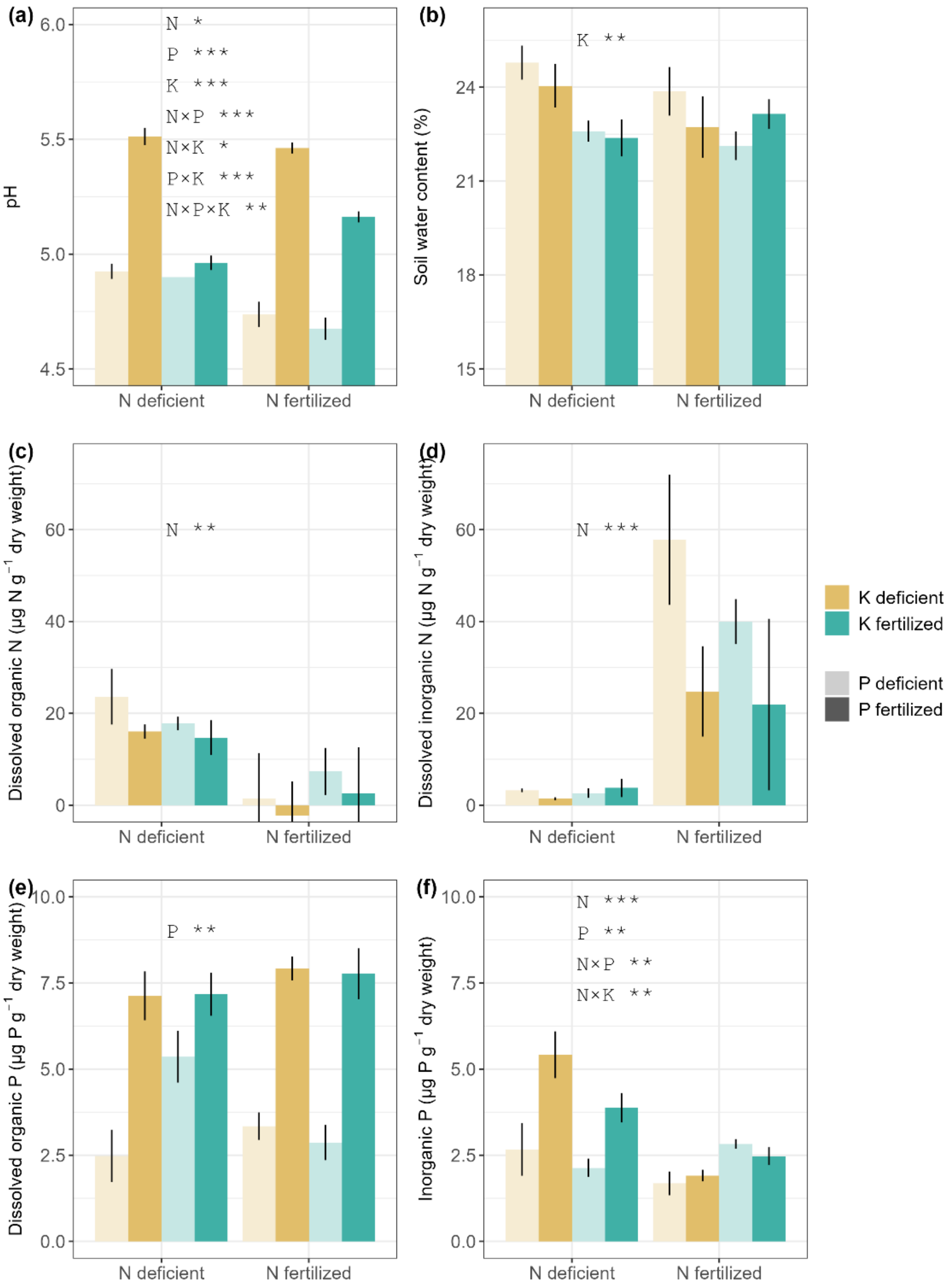

**Fig. S3: The impact of long-term nutrient deficiencies on soil properties and nutrient content.** The y-axes in the boxplots represent soil pH (a), soil water content (b), dissolved organic N (c), dissolved inorganic N (i.e. the sum of $NH_4$ and $NO_3$, d), dissolved organic P (e) and dissolved inorganic P (f). X-axes depict N treatments (left bars: N-deficient, right bars: N-fertilized). Colours indicate K treatments (ochre: K-deficient, cyan: K-fertilized). Colour transparency indicates P treatments (light colors: P-deficient, dark colors: P-fertilized). The bars represent means ± standard error of the mean (n=4). Statistical significance from the three-way ANOVA is reported within each panel: *: $P < 0.05$, **: $P < 0.01$, ***: $P < 0.001$ (See Table **S1**).

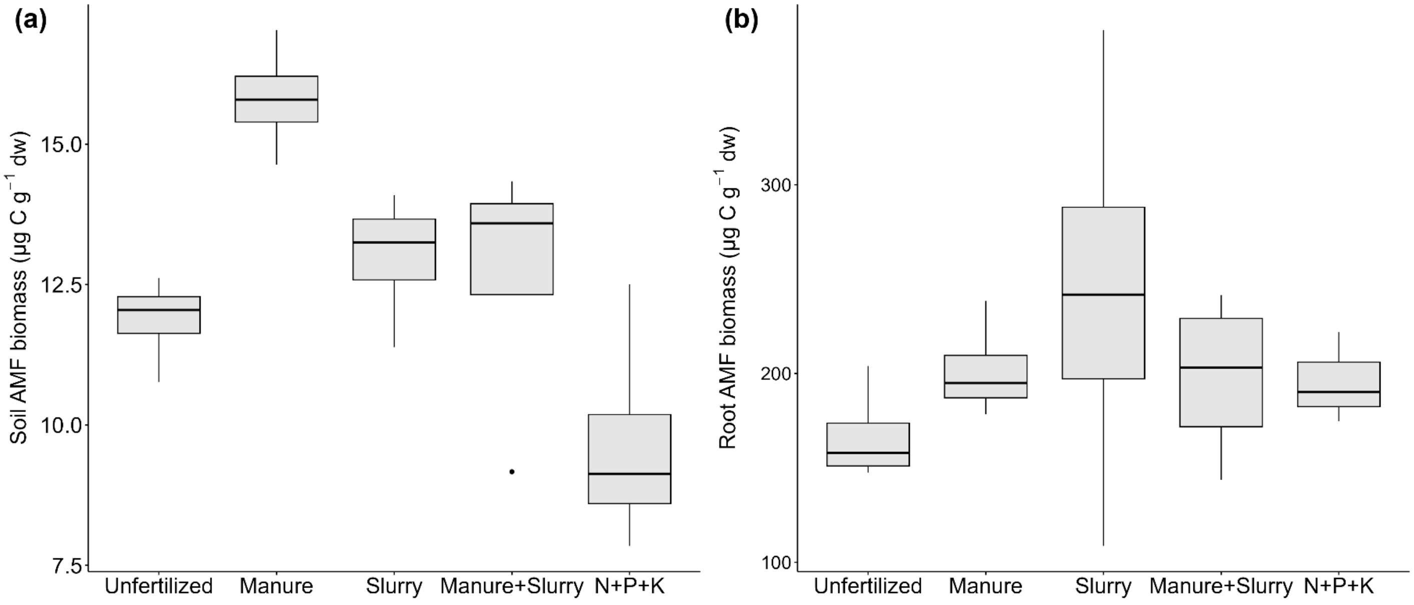


**Fig. S4: Effects of long-term organic fertilization (Manure and slurry) on soil and root AMF biomass.** Organic fertilization increased AMF biomass significantly in soil compared to fully fertilization with NPK, but it had a minor positive effect on root AMF biomass. The y-axes in the boxplot shows a) soil and b) root AMF biomass quantified using 16:1ω5 fatty acid in neutral lipid fraction. The X-axis is associated with the different organic treatments (manure and slurry, and their combination), as well as unfertilized and NPK treatments. Boxplots represent the first and third quartiles and the solid lines represent the median (n=4).

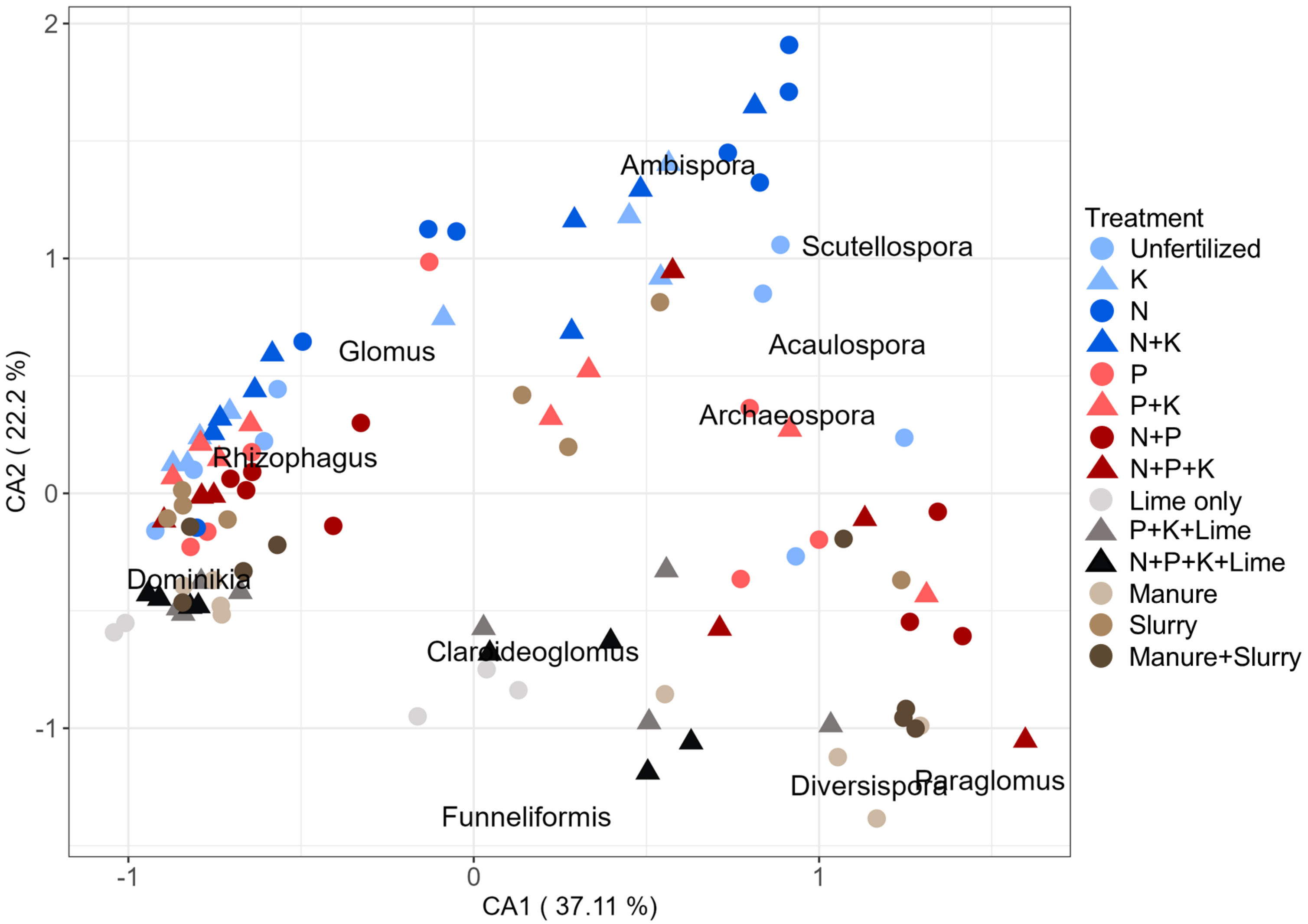


**Fig. S5: Compositions of AMF communities in combined soil and root samples exposed to different long-term nutrient deficiencies evaluated by correspondence analysis (CA).** Lime and organic treatments have different community compositions in comparison with inorganic treatments. Slurry treatment was the only exception, as its community composition is similar to inorganic treatments. The biplot shows correspondence analysis in inorganic, lime and organic treatments in combined soil and root samples. Concerning inorganic treatment, the blue and red colors are indicators of P deficient and fertilized treatments, sequentially. The shapes illustrate K deficient (●) and fertilized (▲) plots. The lightness and darkness of blue and red colors explain N deficient and fertilized treatments, respectively. Other colors are related to lime and organic treatments. the text labels depict AMF taxon at the genus level. Soil and root AMF community compositions were assessed by RNA and DNA amplicon sequencing of 18S rRNA gene, respectively. Root and soil samples can be distinguished in the biplot based on their positioning (Figure **2**).

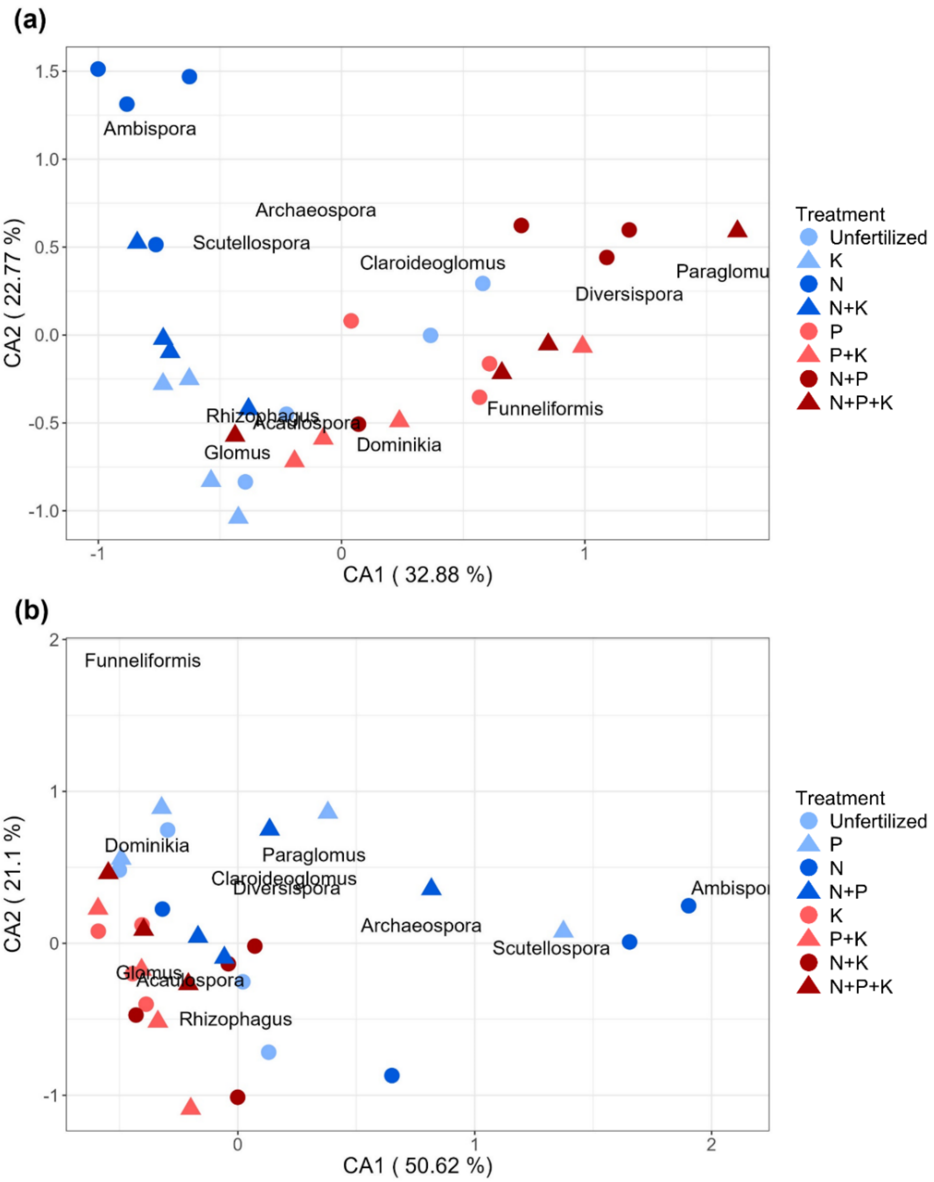


**Fig. S6: The effects of long-term nutrient deficiencies on soil and root AMF community composition based on correspondence analysis (CA).** The soil AMF community composition was significantly affected by N and P treatments (PERMANOVA for P deficient, N deficient and P plus N fertilized treatments: $F$=6.53, $P$<0. 001), whereas the root AMF community composition was influenced by K (PERMANOVA for K deficient and K fertilized treatments: $F$=5.23, $P$<0. 01). The biplot illustrates only the effects of inorganic treatments on soil (a) and root (b) AMF community composition and it is calculated based on CA. In soil samples (a), the blue and red colors indicate P deficient and fertilized plots, respectively, whereas, in roots (b), The blue and red colors show K deficient and fertilized plots, respectively. The shapes show K deficient (●) and fertilized (▲) treatments in soil samples (a) and P deficient (●) and fertilized (▲) in roots (b). Light and dark colors are related to N deficient and fertilized treatments, sequentially, in both samples and the text labels are indicators of AMF genera.

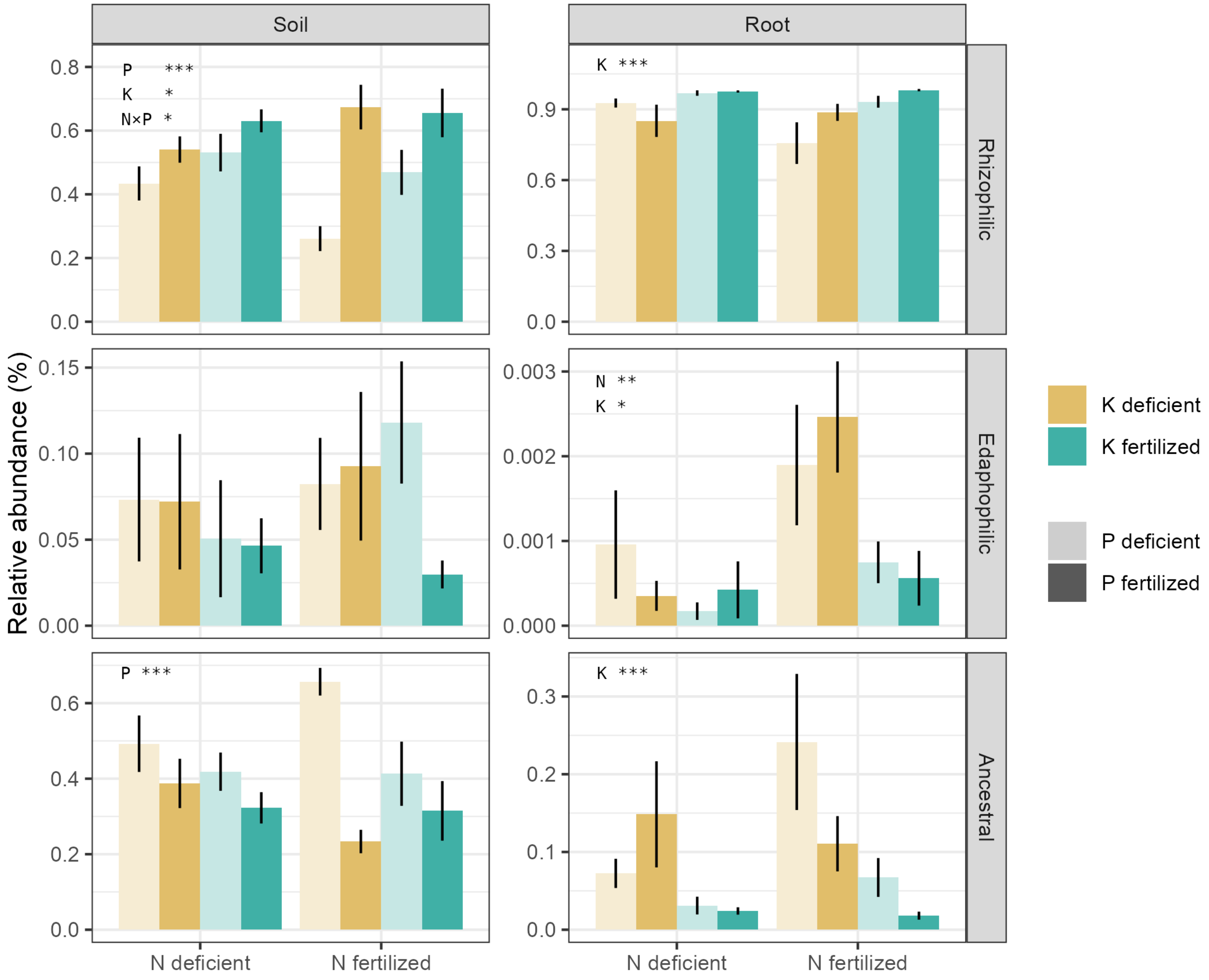


**Fig. S7: The effects of long-term deficiencies of N, P and K on soil and root AMF functional guilds.** Long-term nutrient deficiencies affected different guilds in soil and root samples; however, the responses of different AMF guilds to nutrient deficiencies were different. The panels in columns represent soil and root AMF guilds, and the panels in rows show different types of AMF guilds. The y-axis in each barplot indicates the relative abundance of AMF guilds and the x-axis shows N deficiency (left bars) and N fertilization (right bars). The ochre (deficient) and cyan (fertilized) colors are representative of K in each panel. The alpha shading depicts P deficiency (light colors) and P fertilization (dark colors). The bars indicate means ± standard error of the mean (n=4). Statistical significance from the three-way ANOVA is stated within each panel: *: $P < 0.05$, **: $P < 0.01$, ***: $P < 0.001$ (See Table **1**).

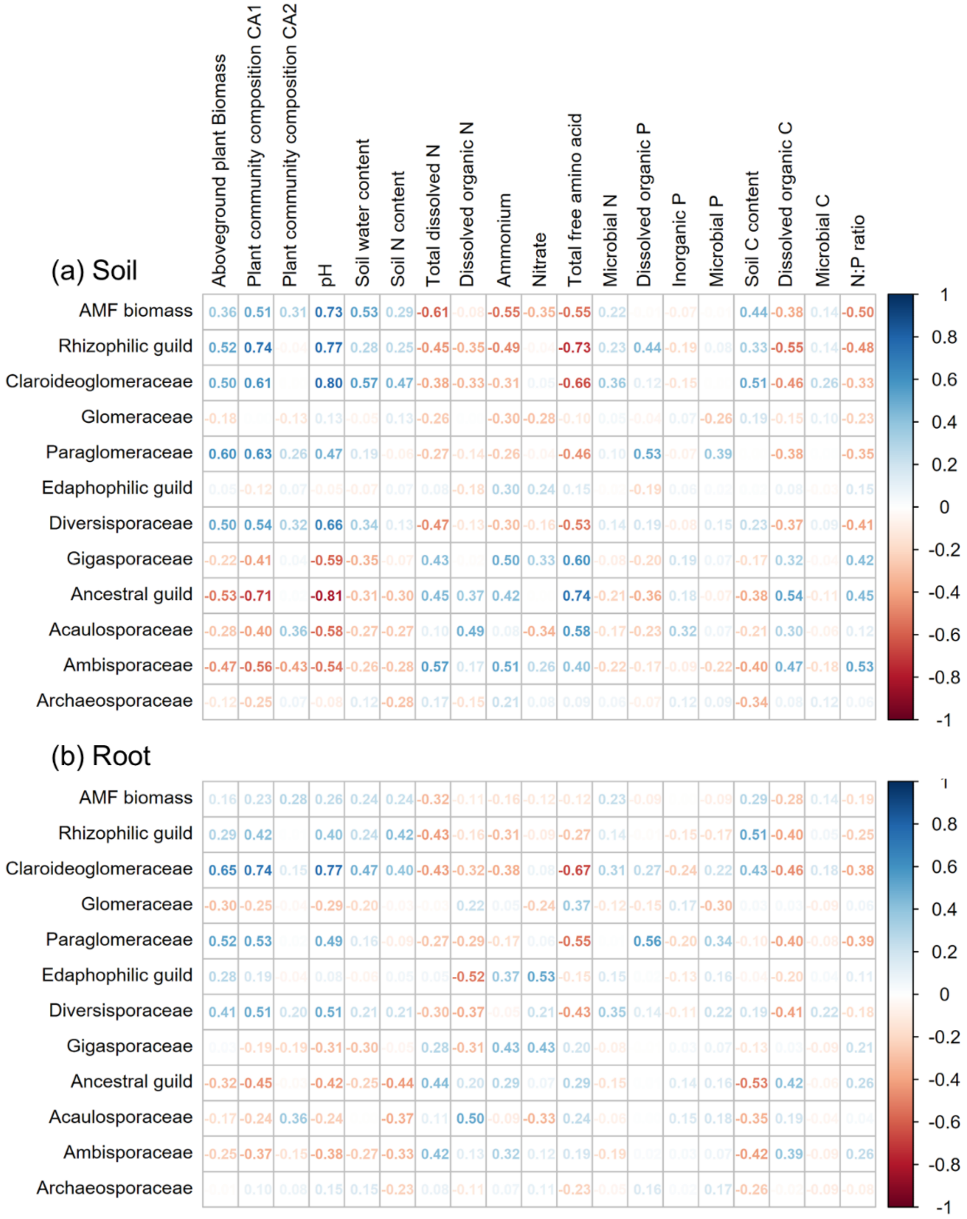


**Fig. S8: Relationship between AMF and environmental parameters across all treatments.** We highlight the strong positive effect of pH on AMF, in contrast to the negative effect of different forms of nitrogen. the correlation plot explains the correlation of soil and root AMF biomass and community composition with plant biomass and community composition, as well as soil parameters considering all treatments (inorganic, organic and lime treatments). The values depicts the correlation coeeficients based on Spearman's correlation and Blue and red colors indicate positive and negative correlations, respectively.

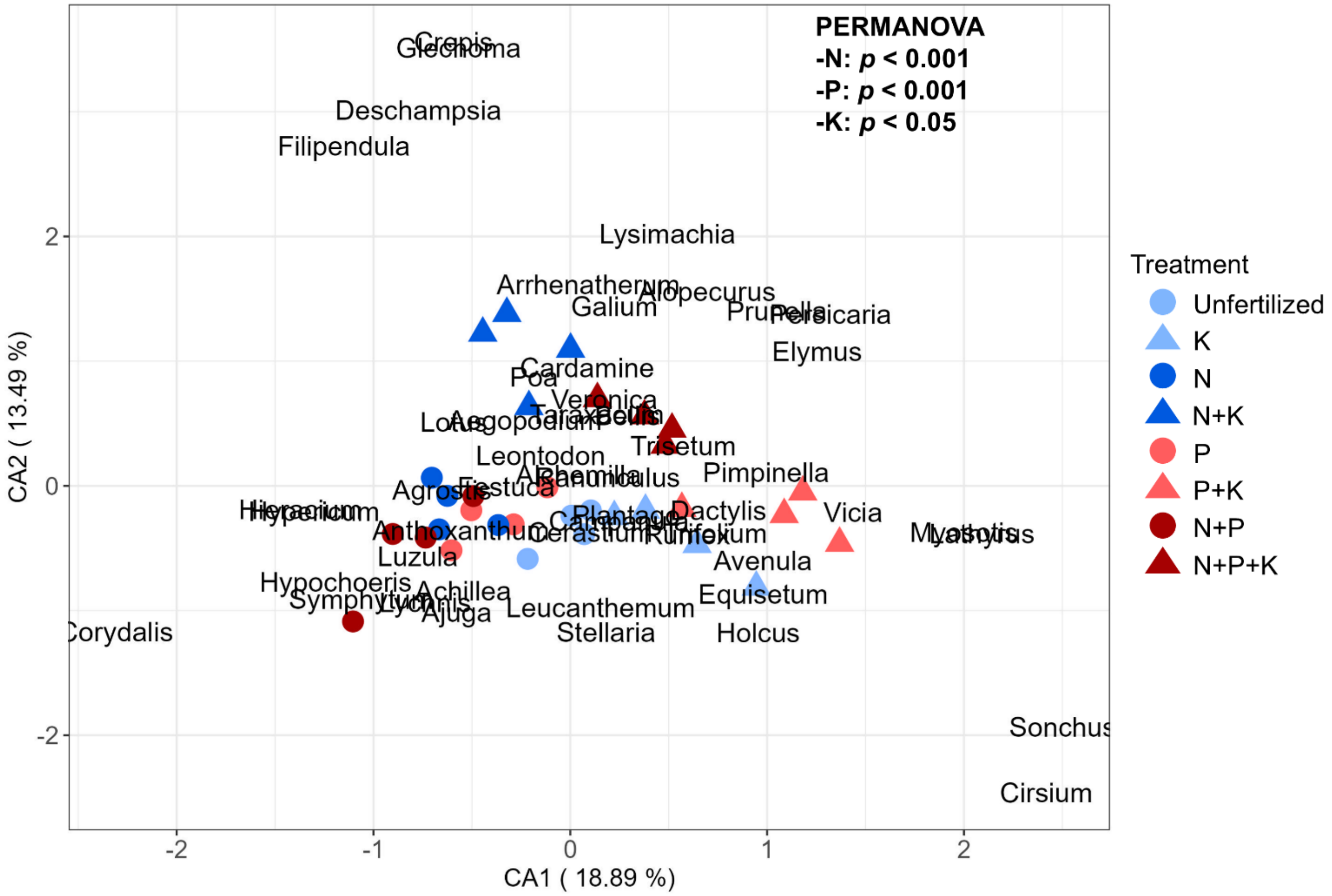


**Fig. S9: The impacts of long-term nutrient deficiencies on plant community composition.** N and P influenced plant community composition (PERMANOVA for P deficient, N deficient and P plus N fertilized treatments : *F*=7.06, *P*<0. 001). The biplot illustrates correspondence analysis of plant community composition in response to long-term deficiencies of N, P and K (inorganic treatments). The blue and red colors show P deficient and fertilized plots, respectively. The shapes show K deficiency (●) and fertilization (▲). Light and dark colors are indicators of N deficient and fertilized treatments, sequentially, and the texts represent plant taxon at the genus level.

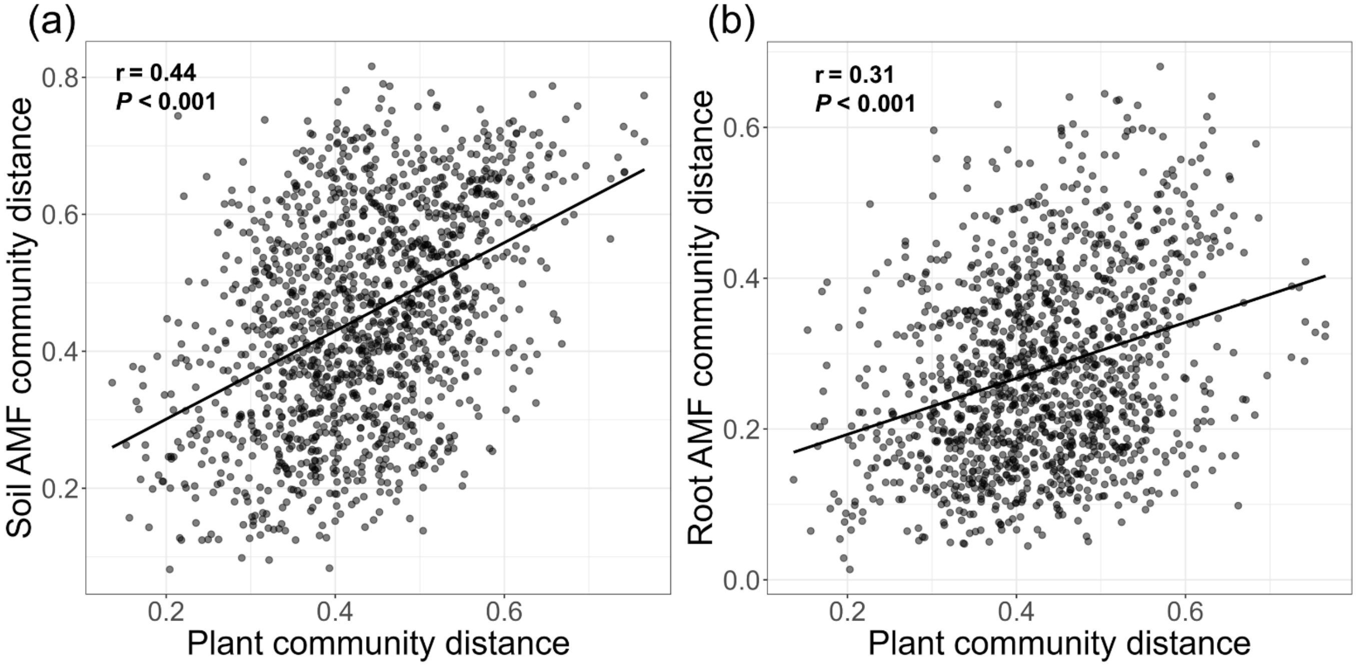


**Fig. S10: Relationship of plant communities with soil and root AMF communities across all treatments.** Mantel test results show correlations between a) plant and soil AMF communities, and b) plant and root-associated AMF community compositions across the inorganic, lime and organic treatments. Each dot indicates a pairwise comparison of community dissimilarities (Bray-Curtis distances) between plots. r shows Pearson's correlation coefficient between the distance matrices.

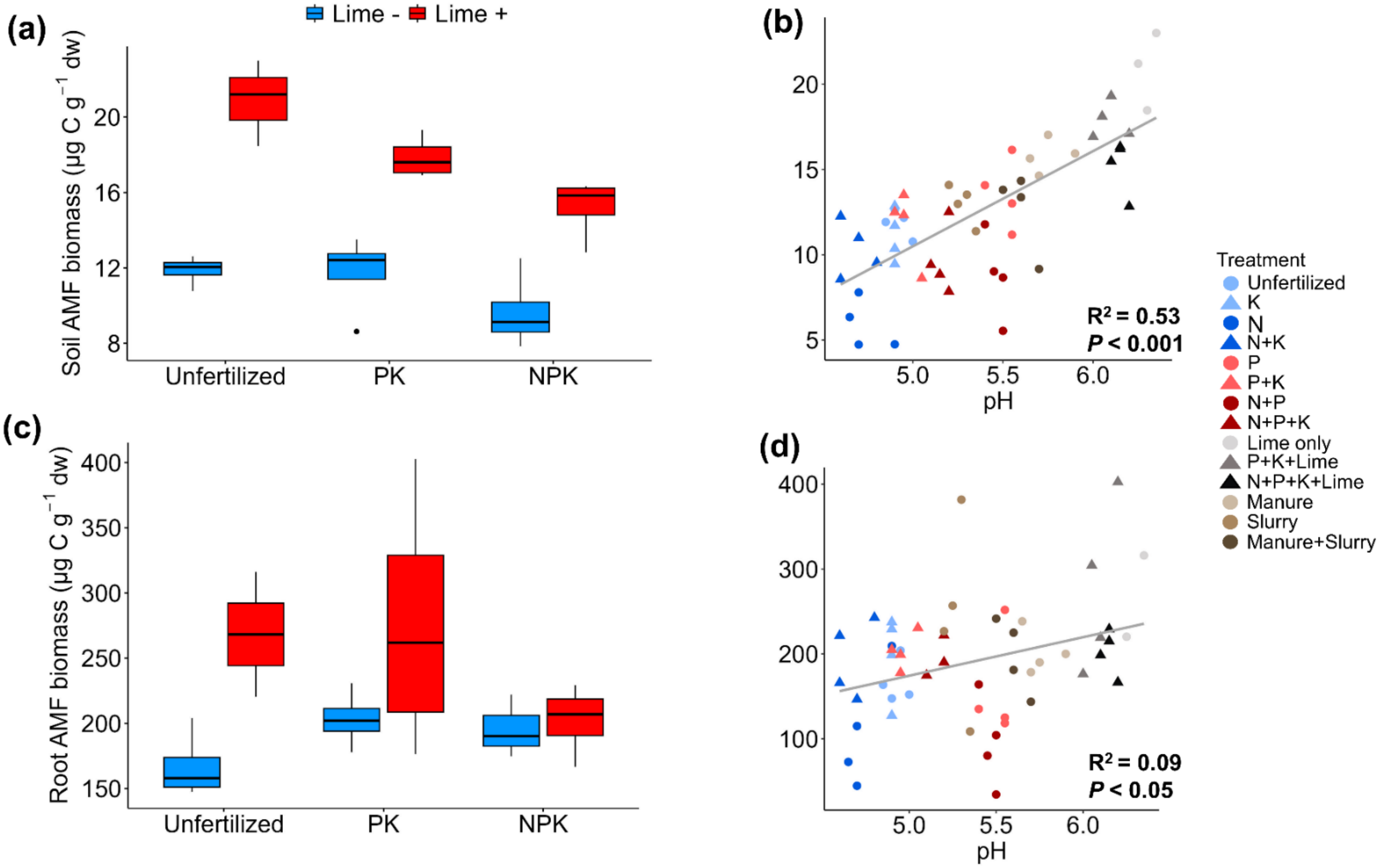


**Fig. S11: The impact of long-term lime application and fertilization regime on soil and root AMF biomass as well as the relationship of soil and root AMF biomass with pH.** Lime significantly increased AMF biomass but had stronger a stronger impact on Soil AMF biomass compared to root AMF biomass. Furthermore, the correlation between pH and soil AMF biomass was stronger Relative to root AMF biomass. The y-axes in boxplots represent a) soil and c) root AMF biomass measured using 16:1 ω 5 fatty acid in neutral lipid fraction. Boxplots indicate the first and third quartiles and the solid lines represent the median (n=4) and the colors are associcated with presence (red) and absence (blue) of Lime. X-axes in boxplots indicate fertilization regiment. The scatterplots illustrate the correlation of b) soil and d) root AMF biomass with pH. In scatterplots, the blue and red colors represent P deficient and fertilized plots, respectively. The shapes indicate K deficient (●) and fertilized (▲) plots. Light and dark colors illustrate N deficient and fertilized treatments, respectively.

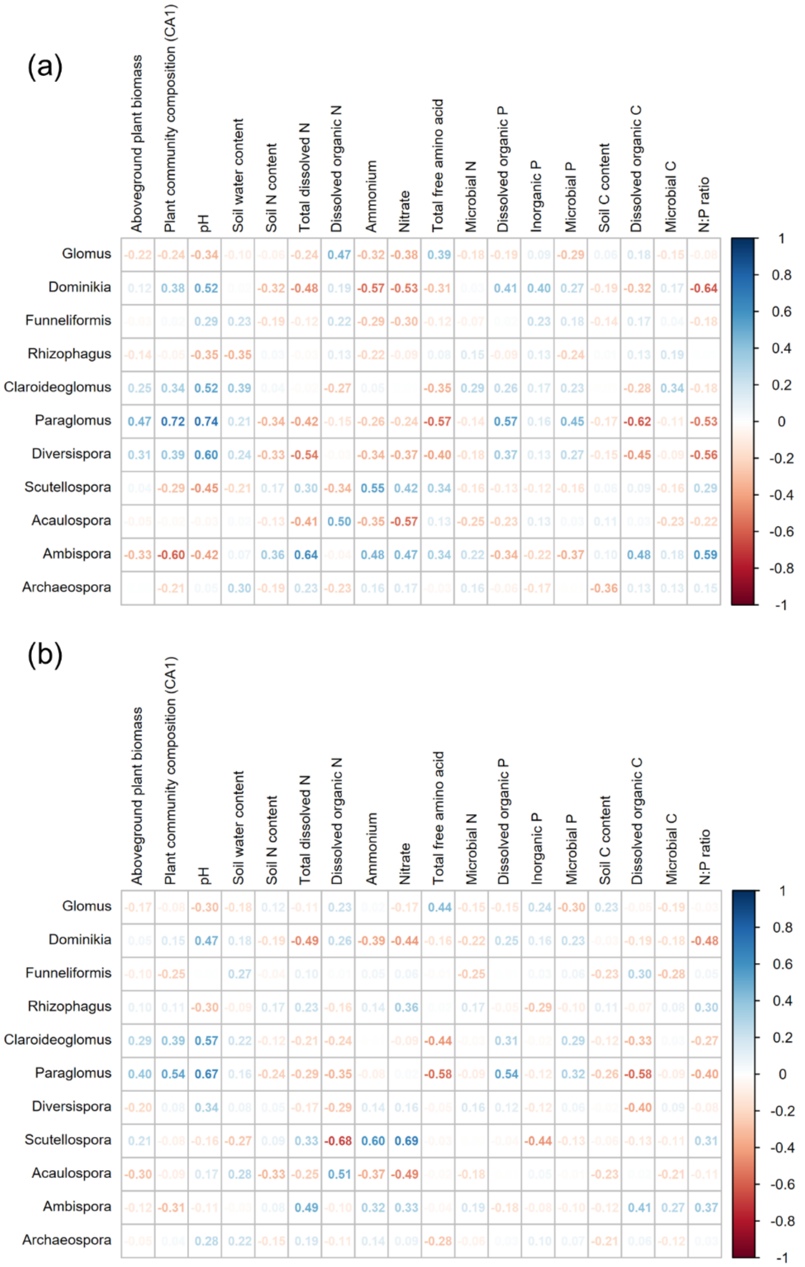


**Fig. S12: Correlations between AMF genera and environmental factors.** Both plant and environmental factors had significant correlations with different AMF genera in soil and root samples. The correlation plot illustrates the relationship between soil (a) and root (b) AMF genera and soil parameters as well as plant biomass and community composition. We considered the correlation in only inorganic treatments and the values show Spearman's correlation coefficient and colors illustrate positive (blue) and negative (red) correlations, respectively.

**Table. S1: Three-way ANOVA results regarding the effect of long-term deficiencies of N, P and K on edaphic factors in inorganic treatments.**

| | -N | -P | -K | -N*-P | -N*-K | -P*-K | -N*-P*-K |
|---|---|---|---|---|---|---|---|
| **pH** | **0.015*↑** | **0.000***↓** | **0.000***↑** | **0.000*** ** | **0.043*** | **0.000*** ** | **0.008** ** |
| **Soil water content** | 0.293 | 0.552 | **0.008**↑** | 0.648 | 0.172 | 0.149 | 0.371 |
| **Soil N content** | 0.055 · ↓ | **0.009**↑** | 0.571 | 0.649 | 0.184 | 0.961 | 0.702 |
| **Total dissolved N** | **0.000***↓** | **0.000***↑** | 0.285 | **0.002** ** | 0.756 | 0.137 | 0.648 |
| **Dissolved organic N** | **0.002**↑** | 0.308 | 0.840 | 0.905 | 0.334 | 0.851 | 0.766 |
| **Dissolved inorganic N** | **0.000***↓** | 0.058 · ↑ | 0.474 | 0.055 · | 0.397 | 0.497 | 0.650 |
| **Ammonium** | **0.000***↓** | **0.008**↑** | 0.238 | **0.009** ** | 0.289 | 0.798 | 0.750 |
| **Nitrate** | **0.000***↓** | 0.162 | 0.157 | 0.173 | 0.125 | 0.428 | 0.466 |
| **Total free amino acids** | 0.798 | **0.000***↑** | 0.075 · ↓ | 0.118 | 0.493 | 0.559 | 0.111 |
| **Microbial N** | 0.191 | 0.112 | 0.722 | 0.856 | 0.697 | 0.616 | 0.407 |
| **Dissolved organic P** | 0.894 | **0.000***↓** | 0.205 | 0.099 · | 0.055 · | 0.167 | 0.086 · |
| **Inorganic P** | **0.000***↑** | **0.002**↓** | 0.770 | **0.001** ** | **0.005** ** | 0.213 | 0.739 |
| **Microbial P** | 0.314 | **0.036*↓** | 0.818 | 0.321 | 0.587 | 0.604 | 0.338 |
| **Soil C content** | 0.020 | **0.031*↑** | 0.065 · ↓ | 0.939 | **0.005** ** | 0.541 | 0.208 |
| **Dissolved organic C** | 0.738 | **0.000***↑** | 0.255 | **0.005** ** | 0.976 | 0.984 | 0.708 |
| **Microbial C** | 0.505 | 0.307 | 0.888 | 0.813 | 0.518 | 0.685 | 0.449 |
| **Plant biomass** | **0.001**↓** | **0.000***↓** | 0.288 | 0.503 | **0.012*** | **0.013*** | 0.428 |

*: $P < 0.05$, **: $P < 0.01$, ***: $P < 0.001$, . : $P < 0.1$, ↑ : positive main effect, ↓ : negative main effect

Shown are P values from a three-way ANOVA using N, P and K deficiency (-N,-P,-K) as factors, including testing their interactions (-N*-P, -N*-K,-P*-K,-N*-P*-K) . The analysis is based on a fully factorial design of N, P and K deficiency with 4 replicate plots per treatment (n=4, total nr of plots: 32). Bold values show *P* below 0.05 and the nutrients in header represent deficiency

**Table. S2: Results of canonical correspondence analysis (CCA) testing the effect of environmental variables on soil and root AMF, and plant community compositions across the inorganic treatments.** Significance of each variable was assessed using permutation tests (n= 9999).

| Environmental factors | df | χ2 | *F*-value | *P*-value |
|---|---|---|---|---|
| **Soil AMF** | | | | |
| **pH** | **1** | **0.1425** | **7.54** | **0.000*****|
| **Dissolved inorganic N** | **1** | **0.0869** | **4.6** | **0.000*****|
| **Plant community composition CA1** | **1** | **0.0568** | **3** | **0.009****|
| Soil water content | 1 | 0.0319 | 1.69 | 0.125 |
| Plant community composition CA2 | 1 | 0.0242 | 1.28 | 0.255 |
| Dissolved organic C | 1 | 0.0217 | 1.15 | 0.329 |
| **Root AMF** | | | | |
| **Dissolved inorganic N** | **1** | **0.0729** | **9.08** | **0.000*****|
| **pH** | **1** | **0.0331** | **4.11** | **0.009****|
| Total free amino acid | 1 | 0.0174 | 2.17 | 0.086 . |
| Plant community composition CA1 | 1 | 0.0136 | 1.7 | 0.156 |
| Inorganic P | 1 | 0.0084 | 1.05 | 0.373 |
| Dissolved organic C | 1 | 0.0069 | 0.85 | 0.471 |
| **Plant** | | | | |
| **pH** | **1** | **0.078** | **2.59** | **0.000*****|
| **Dissolved inorganic N** | **1** | **0.0704** | **2.33** | **0.000*****|
| **Inorganic P** | **1** | **0.0605** | **2.01** | **0.004****|
| **Dissolved organic P** | **1** | **0.0483** | **1.6** | **0.031***|
| Dissolved organic C | 1 | 0.042 | 1.39 | 0.094 . |
| Soil water content | 1 | 0.0412 | 1.37 | 0.102 |

*: $P < 0.05$, **: $P < 0.01$, ***: $P < 0.001$, . : $P < 0.1$

Environmental variables with a Variance Inflation Factor (VIF) less than 10 were selected to avoid the effects of multicollinearity. Tested variables were pH, dissolved inorganic N, dissolved organic P, inorganic P, soil water content, total free amino acids, soil N content, dissolved organic C, plant community composition CA1, CA2, and above ground plant biomass. Plant community composition CA1, CA2, and above ground plant biomass were not tested for plant community composition. Variables with a VIF > 10 as total dissolved N, dissolved organic N, soil C content, N:P ratio, and microbial C, N, and P were filtered. Shown are the six most significant factors for each community, bold environmental factors show *P* below 0.05.

**Table. S3: Results of canonical correspondence analysis (CCA) testing the effect of environmental variables on soil and root AMF, and plant community compositions across all treatments.** Significance of each variable was assessed using permutation tests (n= 9999).

| Environmental factors | df | χ2 | *F*-value | *P*-value |
|---|---|---|---|---|
| **Soil AMF** | | | | |
| **pH** | **1** | **0.2222** | **26.16** | **0.000*****|
| **Dissolved inorganic N** | **1** | **0.0677** | **7.98** | **0.000***** |
| **Plant community composition CA1** | **1** | **0.0361** | **4.25** | **0.001**** |
| **Dissolved organic N** | **1** | **0.0349** | **4.11** | **0.002**** |
| **Dissolved organic P** | **1** | **0.0266** | **3.14** | **0.011*** |
| **Aboveground plan biomass** | **1** | **0.0195** | **2.3** | **0.046*** |
| **Root AMF** | | | | |
| **pH** | **1** | **0.1136** | **27.09** | **0.000***** |
| **Dissolved inorganic N** | **1** | **0.0518** | **12.35** | **0.000***** |
| **Total free amino acid** | **1** | **0.0177** | **4.23** | **0.008**** |
| **Soil water content** | 1 | 0.0082 | 1.96 | 0.111 |
| **Dissolved organic N** | 1 | 0.0077 | 1.84 | 0.127 |
| **Plant community composition CA1** | 1 | 0.0073 | 1.74 | 0.145 |
| **Plant** | | | | |
| **pH** | **1** | **0.1509** | **7.22** | **0.000***** |
| **Soil N content** | **1** | **0.0459** | **2.2** | **0.005**** |
| **Dissolved inorganic N** | **1** | **0.042** | **2.01** | **0.011*** |
| **Inorganic P** | **1** | **0.0357** | **1.71** | **0.032*** |
| **Dissolved organic P** | 1 | 0.0324 | 1.55 | 0.055 . |
| **Dissolved organic N** | 1 | 0.0315 | 1.51 | 0.07 . |

*: $P < 0.05$, **: $P < 0.01$, ***: $P < 0.001$, . : $P < 0.1$

Environmental variables with a Variance Inflation Factor (VIF) less than 10 were selected to avoid the effects of multicollinearity. Tested variables were pH, dissolved inorganic N, dissolved organic N, dissolved organic P, inorganic P, soil water content, total free amino acids, soil N content, dissolved organic C, plant community composition CA1, CA2. Plant community composition CA1, CA2, and above ground plant biomass were not tested for plant community composition. Variables with a VIF > 10 as total dissolved nitrogen, soil C content, N:P ratio, and microbial C, N, and P were filtered. Shown are the six most significant factors for each community, bold environmental factors show *P* below 0.05.
